# Type I interferons promote germinal centers through B cell intrinsic signaling and dendritic cell dependent Th1 and Tfh cell lineages

**DOI:** 10.1101/2020.11.20.390625

**Authors:** Madelene W. Dahlgren, Adam W. Plumb, Kristoffer Niss, Katharina Lahl, Søren Brunak, Bengt Johansson-Lindbom

## Abstract

Type I interferons (IFNs) play an essential role in antiviral immunity, correlate with severity of systemic autoimmune disease, and are likely to represent a key component of mRNA vaccine-adjuvanticity. Relevant to all, type I IFNs can enhance germinal center (GC) B cell responses but underlying signaling pathways are incompletely understood due to pleiotropic effects in multiple cell types. Here, we demonstrate that a succinct type I IFN response promotes GC formation and associated IgG subclass distribution primarily through signaling in cDCs and B cells. Type I IFN signaling in cDCs, distinct from cDC1, stimulates development of separable Tfh and Th1 cell subsets. However, Th cell-derived IFN-γ induces T-bet expression and IgG2c isotype switching prior to this bifurcation and has no evident effects once GCs and *bona fide* Tfh cells developed. This pathway acts in synergy with early B cell-intrinsic type I IFN signaling, which reinforces T-bet expression in B cells and leads to a selective amplification of the IgG2c^+^ GC B cell response. Despite the strong Th1 polarizing effect of type I IFNs, the Tfh cell subset develops into IL-4 producing cells that control the overall magnitude of the GCs and promote generation of IgG1 ^+^ GC B cells. Thus, type I IFNs act on B cells and cDCs to drive GC formation and to coordinate IgG subclass distribution through parallel Th1 and Tfh cell-dependent pathways.

## Introduction

Neutralizing antibody responses develop within germinal centers (GCs); histological structures that arise within lymphoid tissues due to antigen-driven clonal B cell expansion. GCs are also thought to represent key sites for breach of tolerance in development of autoimmune disease *(1)*. GC B cell responses rely on T follicular helper (Tfh) cells, which support B cell expansion, facilitate antibody affinity maturation and eventually select GC B cells into the compartments of long-lived plasma cells or memory B cells *(2)*.

Type I interferons (IFNs), including a single IFN-β and several IFN-a proteins, are rapidly produced in response to viral and bacterial infections and signal through the common and ubiquitously expressed heterodimeric IFN-a receptor (IFNAR) *(3)*. Initially discovered for their ability to induce the “antiviral state” in host cells, type I IFNs are also associated with a plethora of immune-regulatory functions essential for antiviral immunity *(4)*. The importance of type I IFNs in controlling SARS-CoV-2 and their role in preventing severe COVID-19 disease was also recently demonstrated *(5, 6)*. Furthermore, elevated type I IFN production is a hallmark of systemic autoimmune diseases, with a strong correlation between the IFN gene expression signature and clinical manifestations in SLE *(7–10)*. When co-injected with a protein antigen, IFN-a has sufficient adjuvant activity to induce GC formation *(11)* and GC B cell responses have been shown to depend on type I IFNs both during viral infections *(12)* and in autoimmune models *(13, 14)*. While we and others have shown that type I IFN signaling in cDCs augments generation of Tfh cells *(15, 16)*, direct signaling in B- and T-cells has also been implicated in type I IFN-dependent enhancement of humoral immunity and autoimmunity *(11, 12, 14, 17, 18)*. The relative contribution of these pathways, and how they interact with the type I IFN – cDC – Tfh cell axis to enhance and modulate the GC response, is however unclear. Different IgG subclasses are associated with distinct effector mechanisms. In C57BL/6 mice, complement-fixing IgG2c (IgG2a in the BALB/c strain) is more efficient than other subclasses in neutralizing viruses *(19)* and is also associated with more severe pathology in lupus models *(20)*. The transcription factor T-bet induces class switch recombination (CSR) to IgG2a/c *(21)*. T-bet expression in B cells appears to have effects beyond CSR *(22)* and was recently shown to be required for generation of protective anti-influenza stalk region-specific antibodies *(23)*. Mouse IgG2a/c may hence represent a useful surrogate marker for antibody responses driven by T-bet expressing B cells, with relevance for the human setting. Purified IFN-β can enhance production of all IgG subclasses in mice and a similarly broad and type I IFN-dependent isotype distribution is induced through the adjuvant effects of the synthetic dsRNA analogue polyinosinic-polycytidylic acid (poly I:C) *(24)*. Other studies suggest that type I IFNs preferentially promote IgG2a/c *(25, 26)*, and B cell-intrinsic type I IFN signaling mediates IgG2a/c CSR in the T cell-independent response to NP-Ficoll *(27)*. The importance of this pathway in GC B cell responses is however less clear and the IgG2a/c subclass has more frequently been associated with strong Th1 immunity and IFN-γ production *(28, 29)*.

Type I IFNs can both inhibit and promote generation of IFN-γ producing Th1 cells, with a general trend that sustained type I IFN responses during chronic infections suppress Th1 immunity *(4)*. Through binding to endosomal TLR3 and the cytoplasmic RNA helicase MDA5, poly I:C instead triggers a transient systemic type I IFN response that peaks 3-6 hours after injection *(30, 31)*, and an almost identical pattern in type I IFN production is observed for novel mRNA vaccination regimens *(32)*. In contrast to the chronic infection models, this short-lived type I IFN response leads to Th1-biased immunity with abundant IFN-γ production from CD4 and CD8 T cells and in particular from NK cells *(30)*. How this IFN-γ response develops, and what impact it has on GC B cell differentiation remains to be determined.

In the current study we have identified the pathways by which a succinct type I IFN response induced by poly I:C drives formation of GCs characterized by a broad IgG subclass distribution. Our experiments also reveal the trajectories downstream of type I IFNs that promote T-bet expression and IgG2c CSR in developing GC B cells.

## Results

### Reduced GC B cell response in Ifnar1^-/-^ mice involves loss of both IgG1^+^ and IgG2c^+^ GC B cells

To determine how type I IFNs affect GC B cell differentiation at the level of individual IgG subclasses, *Ifnar1^-/-^* mice were immunized i.p. with OVA and poly I:C. To visualize concurrent CD4 T cell responses (see below), all mice received 5000 CD45 congenic OT-II cells. Thus, all cells are unable to respond to type I IFNs in these recipients except for a small and physiologically relevant number of antigen-specific CD4 T cells. In contrast to the strong splenic GC B cell response detected in wt controls after eight days, immunization of *Ifnar1^-/-^* mice did not lead to an increase in percentage or number of GC B cells (Fig. 1A and B). The percentage and number of GC B cells were however similar between unimmunized wt and *Ifnar1^-/-^* mice (Fig. 1A and B), which is in agreement with reports of an important role for B cell IFN-γ receptor signaling, and not type I IFNs, in driving spontaneous GC formation in wt C57BL/6 mice *(33, 34)*. The response induced by immunization of wt mice involved equal percentages (Fig. 1C and D) and numbers (Fig. 1E) of IgG1^+^ and IgG2c^+^ GC B cells. Due to the robust magnitude of this response, pre-existing GCs made a very small and negligible contribution to the isotypes expressed by GC B cells in immunized wt mice. In *Ifnar1^-/-^* mice, we observed a significantly increased percentage of IgG1 ^+^ GC B cells after immunization, indicating that an IgG1 ^+^ GC B cell response to some extent had developed also in the absence of type I IFN signaling (Fig. 1 C and D). However, in this set of experiments we could not detect a corresponding increase in the number of IgG1 ^+^ GC B cells (Fig. 1E). There was no significant increase in either percentage (Fig. 1C and D) or number (Fig. 1E) of IgG2c^+^ GC B cells in immunized as compared to non-immunized *Ifnar1^-/-^* mice.

**Fig 1.**
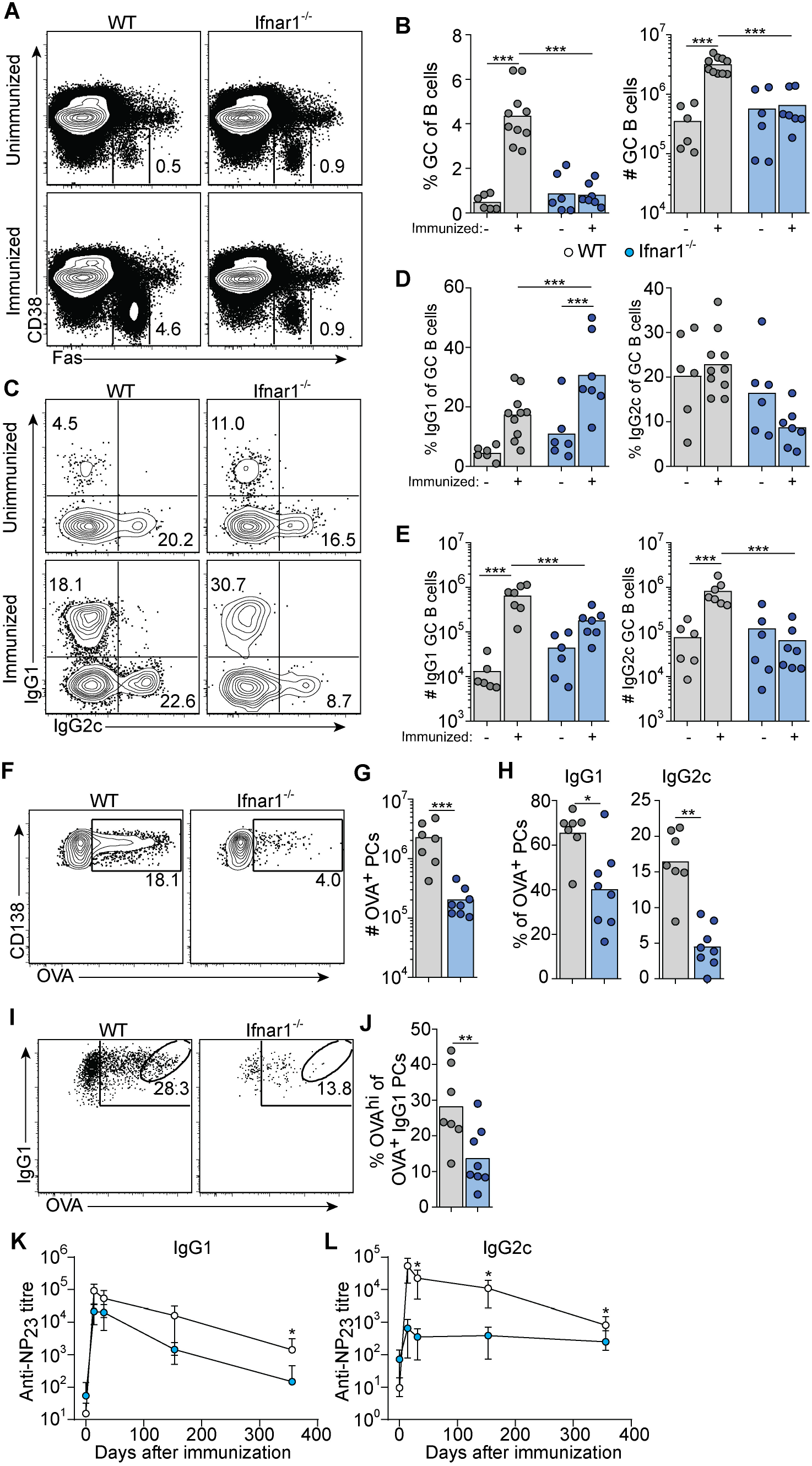
Attenuated primary GC B cell responses in *Ifnar1^-/-^* mice. 5000 OT-II cells were transferred into WT or *Ifnar1^-/-^* mice which were subsequently immunized with OVA **(A-J)** or NP-OVA **(K-L)** + poly I:C. Splenocytes were analyzed 8 dpi **(A-J)**. (A) Flow plots of frequency of GC B cells within total B cells. **(B)** Percentage and numbers of GC B cells. **(C)** Flow plots of IgG1 and IgG2c expression by GC B cells. **(D)** Percentage and **(E)** total numbers of IgG1^+^ and IgG2c^+^ GC B cells. **(F)** Flow plots of intracellular OVA binding of PC. **(G)** Total number of OVA^+^ PCs. **(H)** Percentage of IgG1^+^ and IgG2c^+^ cells in OVA^+^ PCs. **(I)** Flow plots and **(J)** frequency of OVA^hi^ PCs within total OVA^+^ IgG1^+^ PCs. Serum titers of NP_23_-specific **(K)** IgG1 and **(L)** IgG2c, before after immunization. Results are pooled from three **(A-D)**, two **(E-J)** or one **(K-L)** individual experiments, each symbol represents one mouse **(A-J)** or n=6 per group **(K-L)**. **p<0.01 and ***p<0.001.

The majority of antigen-specific plasma cells (PCs) present in spleen around eight days after immunization are thought to be derived from GCs *(35)*. The total number of splenic OVA-binding CD138^+^ PCs was approximately 10-fold lower in *Ifnar1*^-/-^ compared to wt mice (Fig. 1F and G), and both IgG1- and in particular IgG2c-producing cells were affected (Fig. 1H). In addition, the few IgG1^+^ PCs present in *Ifnar1*^-/-^ mice appeared to produce antibodies of lower affinity than their counterparts in wt animals, as indicated by fewer OVA binding IgG1 cells binding high levels of OVA (Fig. 1I and J). These results were further corroborated by serological studies where *Ifnar1^-/-^* mice essentially failed to mount an NP-specific IgG2c response after administration of NP-conjugated OVA plus poly I:C whereas their NP-specific IgG1 titers were reduced only 6.9- and 4.2-fold at two and four weeks after immunization, respectively (Fig. 1K and L). Antigen-specific IgG1 titers however then declined more rapidly in the *Ifnar1^-/-^* mice and after one year they had an almost 80-fold lower NP-specific IgG1 titer than wt controls, indicating that the longevity of the specific IgG1 response was affected in the absence of type I IFN signaling (Fig. 1K). Collectively these results demonstrate that type I IFNs are essential for GC formation induced by poly I:C, affecting both the quantity and quality of the response and, moreover, demonstrate that type I IFN signaling promotes the generation of both IgG1^+^ and, in particular, IgG2c^+^ GC B cells.

### Tfh cells and IFN-γ producing Th1 cells develop concurrently in response to poly I:C and are both reduced in Ifnar1^-/-^ mice

We have previously shown that early Tfh cell fate commitment is reduced in *Ifnar1^-/-^* mice (as assessed three days after immunization) *(15)*. To assess the relationship between Tfh and Th1 cell development, and to determine how type I IFN impacts on the accumulation of these subsets at the peak of the GC reaction, we analyzed transferred OT-II cells in the same cohorts of mice as described above. *Ifnar1*^-/-^ mice had fewer total OT-II cells in their spleens as compared to wt controls, confirming that type I IFNs enhance expansion and survival of activated CD4^+^ T cells *(36, 37)* (Fig. 2A). Furthermore, Tfh cells were additionally affected in the *Ifnar1^-/-^* mice, as evident from a significantly reduced percentage of CXCR5^high^ Bcl6^+^ OT-II cells (Fig. 2B and C). T-bet drives Th1 cell differentiation and is required for IFN-γ expression by CD4^+^ T cells *(38)*. OT-II cells with detectable T-bet expression were confined to the Bcl6^-^ subset in the wt animals and in agreement with previously published results *(30)* IFN-γ expression was essentially lost in the *Ifnar1*^-/-^ recipients, as was expression of T-bet (Fig. 2D-G). Collectively, these results show that Tfh cells and IFN-γ producing Th1 cells exist as mutually exclusive subsets at the peak of the GC reaction and that both subsets depend on type I IFNs for their development.

**Fig 2.**
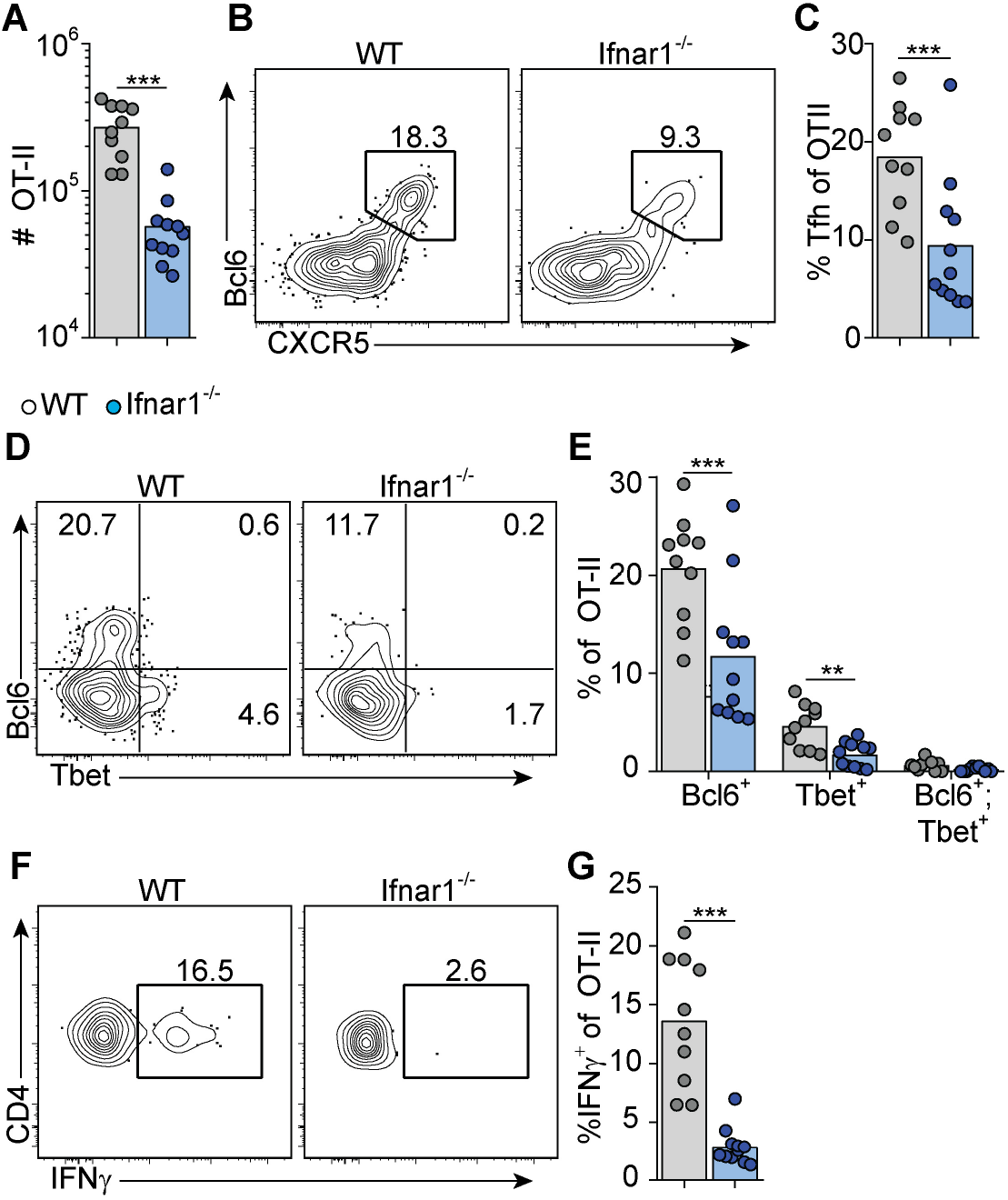
Type I IFN signaling augments Tfh and is required for Th1 cell differentiation. WT and *Ifnar1^-/-^* mice were transferred with 5000 OT-II cells and immunized with OVA + pI:C. Splenocytes were analyzed 8 dpi. **(A)** Number of OT-II cells. **(B)** Flow plots of Bcl6 versus CXCR5 staining in OT-II cells. **(C)** Percentage of Tfh cells. **(D)** Representative plots of Bcl6 and T-bet expression and **(E)** percentage of single- or co-expression of Bcl6 and T-bet by OT-II cells. **(F)** Representative plots and **(G)** percentage of IFN-γ expression by OT-II cells. Results are pooled from three individual experiments and each symbol represents one mouse. **p<0.01 and ***p<0.001.

### Early IFN-γ derived from cognate CD4^+^ T cells acts directly on B cells to drive IgG2c CSR without enhancing the overall magnitude of the GC B cell response

IFN-γ represents a well-established *in vitro* switch factor for the IgG2a/c subclass *(39)*. B cell intrinsic IFN-γ signaling can also underlie GC formation in murine autoimmune models *(33, 34)*. To determine the role of B cell intrinsic IFN-γ signaling in the GC B cell response driven by OVA/poly I:C, we made mixed bone marrow (BM) chimeric mice. Irradiated wt mice were grafted with wt BM (CD45.1^+^, CD45.2^+^) mixed with *Ifngr1^-/-^* or wt control BM (both CD45.2 single positive) at a 1:1 ratio. Analysis of splenocytes eight days after immunization showed that GC B cells developed equally well from *Ifngr1^-/-^* and wt B cells (Fig. 3A and B), demonstrating that IFN-γ signaling in B cells is not involved in type I IFN dependent GC B cell expansion. In contrast, CSR to IgG2c was strongly reduced in the *Ifngr1^-/-^* as compared to wt GC B cells present in the same mouse (Fig. 3A and B). In agreement with the ability of IFN-γ to inhibit CSR to IgG1 and IgE *(28, 39)*, the impaired IgG2c CSR in *Ifngr1^-/-^* B cells was compensated by an increased percentage of IgG1^+^ GC B cells (Fig. 3A and B). Given that IL-27 represents an additional IgG2a/c switch factor *(40)* and can be produced by myeloid cells in response to type I IFNs *(41, 42)*, we performed analogous experiments with mixed *wt/Il27ra^-/-^* BM chimeras. These experiments did however not reveal any significant differences between wt and *Il27ra^-/-^* B cells (Fig. S1A and B).

**Fig 3.**
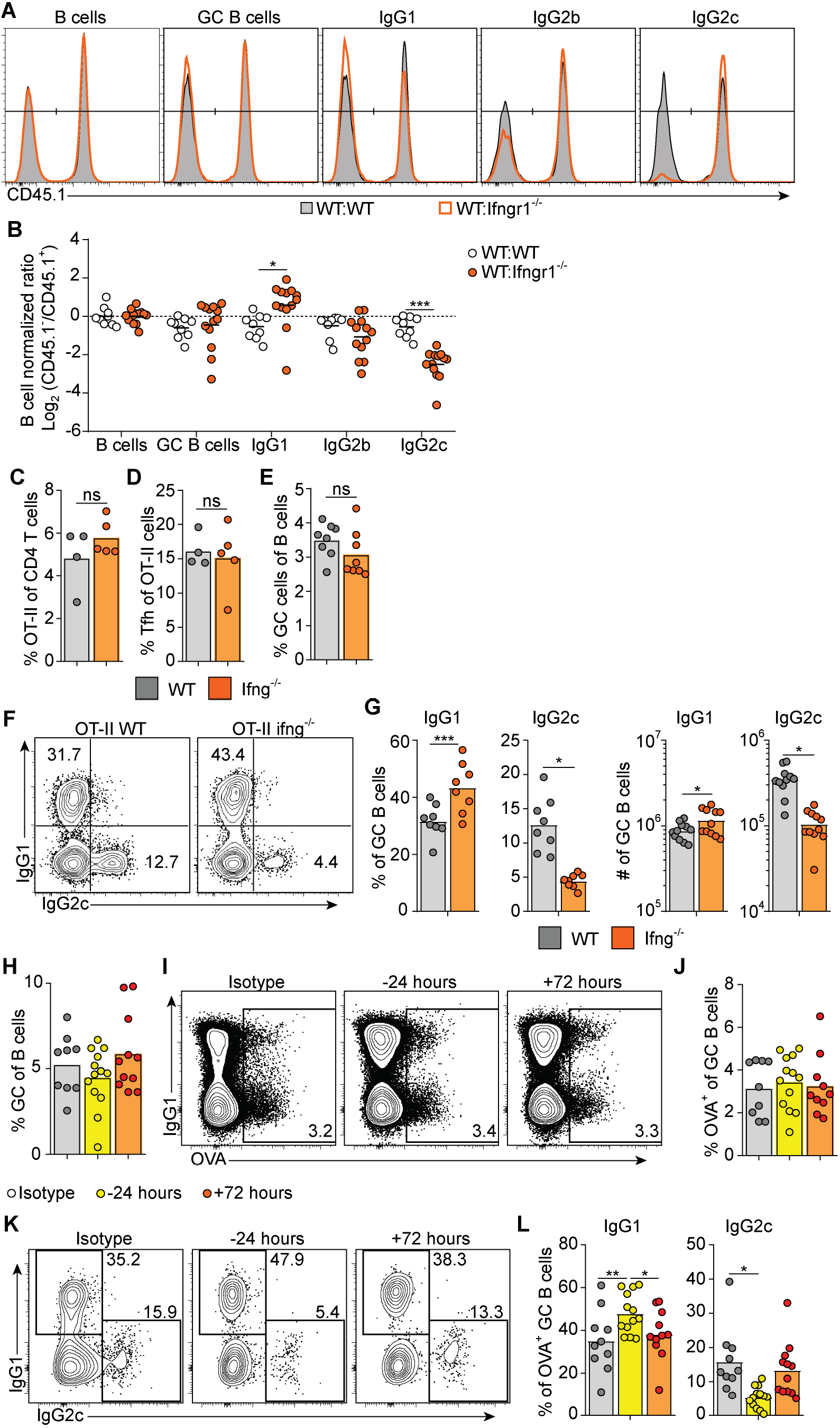
Early IFN-γ derived from cognate CD4 T cells acts directly on B cells to drive IgG2c CSR. **(A-B)** Mixed chimeras were generated by reconstituting WT recipients with a 1:1 mix of congenic WT and WT or *Ifngr1^-/-^* BM cells. 8-10 weeks after reconstitution, chimeras were immunized with OVA + poly I:C, and splenic GC B cell responses were analyzed 8 dpi. **(A)** Histograms and **(B)** log_2_ normalized ratio of WT:WT and WT:*Ifngr1^-/-^* chimeras showing the distribution of B cells, GC B cells and GC B cells expressing IgG isotypes. **(C-G)** *Ifng^-/-^* mice were transferred with 50 000 WT or *Ifng^-/-^* OT-II cells and immunized with OVA + poly I:C. and splenocytes were analyzed 8 days later. Percentage of **(C)** OT-II cells in CD4 T cells, **(D)** Tfh within OT-II cells and **(E)** GC B cells of total B cells. **(F)** Flow plots **(G)** percentage and numbers of IgG1 and IgG2c expressing GC B cells. **(H-L)** WT mice were treated i.p. with 1 mg anti-IFN-γ mAb or isotype control 16 hrs before or 72 hrs after immunization with OVA plus poly I:C. Mice were transferred with 50 000 OT-II 16 hrs before immunization and splenocytes were analyzed 8 dpi. **(H)** Percentage GC B cells of B cells. **(I)** Flow plots and **(J)** percentage of OVA^+^ GC B cells. **(K)** Flow plots and **(L)** percentage of IgG1^+^ and IgG2c^+^ cells in OVA^+^ GC B cells. Pooled results from one **(C-D)** two **(E-G)** or three experiments are shown. Each symbol represents one mouse. ns = not significant, *p≤0.05 and **p<0.01.

To determine if IFN-γ from cognate CD4^+^ T cells is sufficient for driving IgG2c CSR in B cells, we transferred IFN-γ sufficient or deficient OT-II cells into *Ifng^-/-^* mice and again evaluated the splenic response eight days after immunization. Wt and *Ifng^-/-^* OT-II cells had expanded and differentiated into Tfh cells equally efficient in the *Ifng^-/-^* recipient mice (Fig. 3C and D). Additionally, IFN-γ-deficient OT-II cells supported a GC B cell response of a comparable magnitude as their wt counterparts (Fig. 3E). However, compared to mice receiving wt OT-II cells, recipients of *Ifng^-/-^* OT-II cells developed GCs with a strong and selective reduction in IgG2c, again compensated by an increase in IgG1 (Fig. 3F and G). These experiments thus demonstrate an important role for IFN-γ derived from cognate CD4^+^ T cells in stimulating IgG2c CSR, but not for enhancing the overall magnitude of the GCs.

IFN-γ producing Tfh cells have been suggested to underpin IgG2a/c CSR within GCs *(43)*. Given that Tfh cells largely lacked expression of T-bet at the peak of the poly I:C driven GC response (see Fig. 2 D and E), we considered the possibility that IFN-γ stimulates IgG2c CSR at an earlier stage and before evident GC formation. To assess this, we injected an IFN-γ neutralizing antibody either before or 72 hours after immunization. Regardless of the timing, this treatment had no impact on the percentage of total GC B cells detected eight days after immunization and staining with fluorescently labeled OVA confirmed an equal percentage of OVA-specific GC B cells in both experimental groups and in mice receiving an isotype control mAb (Fig. 3H-J). However, treatment with anti-IFN-γ before immunization resulted in significantly fewer OVA-specific GC B cells expressing IgG2c as compared to isotype control treated mice (Fig. 3K and L). This reduction was again reciprocated by an increased percentage of cells expressing IgG1 (Fig. 3K and L). No such effects on the IgG subclass distribution were observed when treatment with anti-IFN-γ instead was started 72 hours after immunization (Fig. 3K and L). Accordingly, IFN-γ acts on B cells within the first 72 hours after immunization to initiate the IgG2c CSR process and thereafter appears to be redundant for the IgG2c associated GC B cell response.

### B cell intrinsic type I IFN signaling acts in synergy with type I IFN dependent IFN-γ to select the IgG2c subclass

The finding that B cell intrinsic IFN-γ signaling involved in IgG2c CSR precedes GC formation predicts that 1) type I IFN dependent priming of IFN-γ producing Th1 cells also occurs early in the response and 2) cognate IFN-γ producing CD4 T cells appear earlier than detectable GCs. To test the first prediction, we neutralized type I IFN signaling by injecting an anti-IFNAR mAb either 16 hours before or 24 hours after immunization. Neutralization of type I IFN signaling before, but not after, immunization inhibited OT-II cell expansion and differentiation into Tfh cells as assessed day eight (Fig. 4A and B). As predicted, IFN-γ production by transferred OT-II cells was completely inhibited at this time point (Fig. 4C). Consistent with the reduced number of Tfh cells, the overall magnitude of the GC B cell response was reduced when anti-IFNAR treatment preceded immunization (Fig. 4D). Additionally, the numbers of IgG2c^+^ GC B cells were reduced after early but not late IFNAR blockade (Fig. 4E). In the same type of neutralization experiment we also confirmed that NP-specific IgG2c was reduced > 10-fold in sera 14 days after immunization when the mice received anti-IFNAR mAb before, but not 24 hours after, immunization (Fig. 4F). The type I IFN dependent signaling event that underlies Th1 development and IFN-γ dependent IgG2c CSR is therefore initiated during the first 24 hours of the poly I:C driven response.

**Fig 4.**
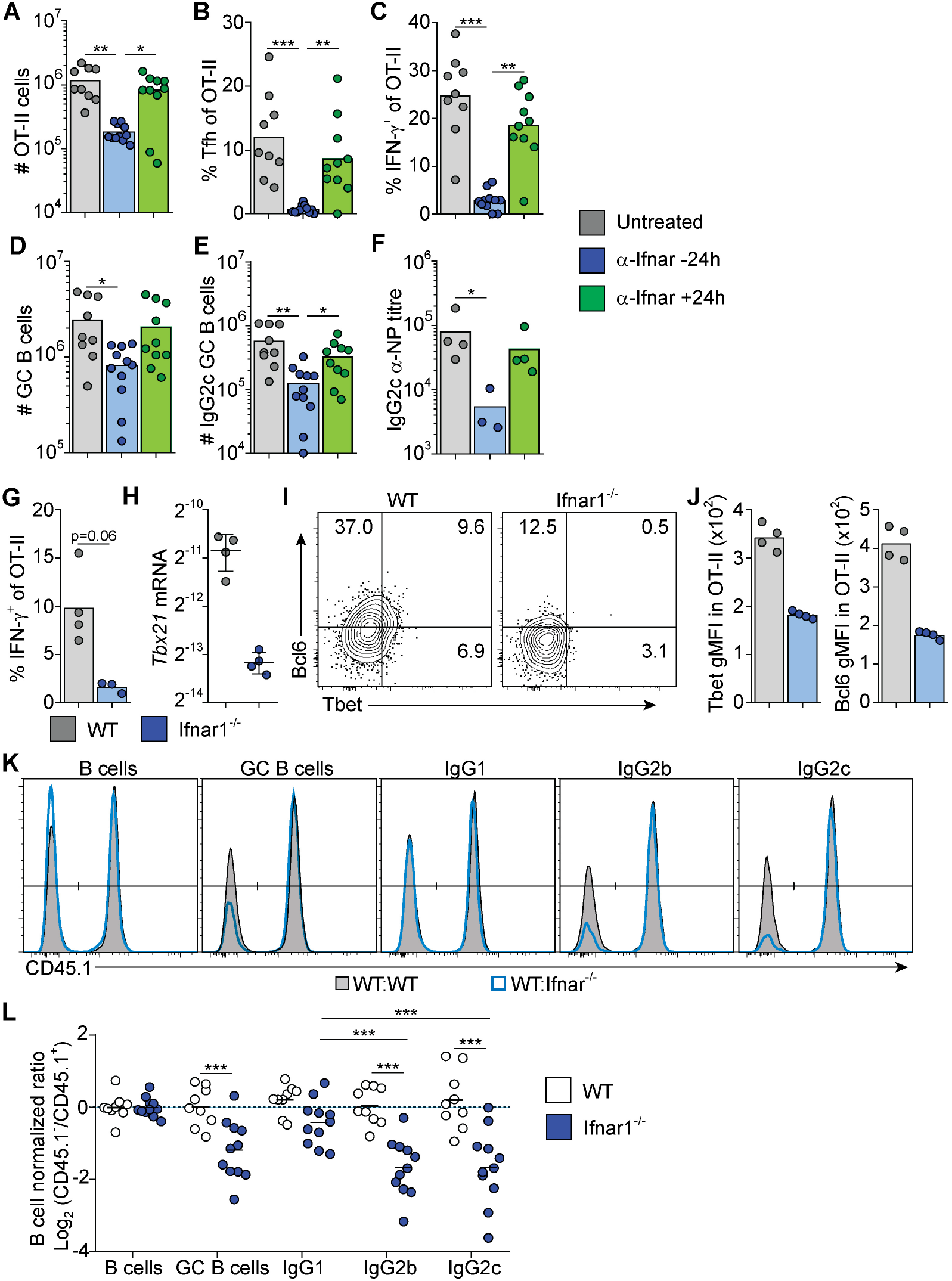
Type I IFN signaling is active within the first 24 hrs and acts directly on B cells. **(A-F)** WT mice were treated i.p. with 1 mg anti-IFNAR mAb or isotype control 16 hrs before or 24 hrs after immunization with **(A-E)** OVA or **(F)** NP-OVA plus pI:C. Mice in **(A-E)** and **(G-J)** were transferred with OT-II 16 h before immunization and splenocytes were analyzed 8 dpi. **(A)** Number of OT-II cells, percentage of **(B)** Tfh and **(C)** IFN-γ^+^ cells of total OT-II cells. **(D)** Number of GC B cells and **(E)** IgG2c GC B cells. **(F)** Serum titers of NP23-specific IgG2c two weeks post-immunization. **(G-J)** WT and *Ifnar1^-/-^* mice were transferred with 500 000 OT-II cells and immunized with OVA + poly I:C. Splenic OT-II cells were sorted 48 hrs postimmunization for qrt-PCR or analyzed by flow cytometry. **(G)** Percentage of total OT-II cells expressing IFN-γ. **(H)** OT-II cell *Tbx21* mRNA levels. **(I)** Representative flow plots and **(J)** gMFI of T-bet and Bcl6 expression by total OT-II cells. **(H-I)** Mixed chimeras were generated by reconstituting WT recipients with a 1:1 mix of congenic WT and WT or *Ifnar1^-/-^* BM cells. 8-10 weeks after reconstitution, chimeras were immunized with OVA + poly I:C, and splenic GC B cell responses were analyzed 8 dpi. **(H)** Histograms and **(I)** log_2_ normalized ratio of WT:WT (shaded) and *WT:Ifnar1^-/-^* chimeras (blue) showing the distribution of B cells, GC B cells and GC B cells expressing indicated IgG isotypes. Pooled results from one **(F-J)** or three experiments. Each symbol represents one mouse. *p≤0.05, **p<0.01 and ***p<0.001.

To confirm that IFN-γ producing Th cells appear within the first 72 hours post-immunization, and to verify their dependence on upstream type I IFN signaling, we analyzed early Th cell differentiation in wt and *Ifnar1^-/-^* mice, respectively. IFN-γ producing OT-II cells were indeed detectable in wt mice 3 days after administration of OVA and poly I:C and the IFN-γ response was reduced in the *Ifnar1*^-/-^ mice also at this earlier time point (Fig. 4G). This correlated with significantly higher *Tbx21* mRNA (Fig. 4H) and T-bet protein expression (Fig. 4 I and J) in OT-II cells recovered from wt compared to *Ifnar1^-/-^* recipients. OT-II cells recovered from wt animals did however not express T-bet and Bcl6 in a mutually exclusive manner indicating the absence of clear Th1 versus Tfh cell dichotomy at this early stage of the response (Fig. 4I). Based on these results we conclude that type I IFNs drive development of the IFN-γ producing CD4 T cells that underpin IgG2c CSR within the first few days after immunization, prior to evident GC formation and appearance of fully committed Th1 and Tfh cells.

To determine how and to what extent direct type I IFN signaling in B cells influences the GC B cell response and IgG subclass selection after immunization with OVA/poly I:C, we again generated mixed BM chimeras, now by reconstituting irradiated wt mice with a 1:1 mix of wt with *Ifnar1^-/-^* or wt (control) BM. Analysis of immunized chimeric mice revealed that *Ifnar^-/-^* B cells contributed to the GC B cell response to a lesser extent than IFNAR sufficient B cells present in the same animal. In this competitive setting, there was an approximately 2-fold lower number of *Ifnar1^-/-^* than wt GC B cells in the spleen (Fig. 4K and L). Strikingly, this reduction was mostly caused by a loss in IgG2c^+^ and IgG2b^+^ GC B cells; the frequency of GC B cells expressing either of these IgG subclasses was approximately 2-fold lower in the *Ifnar1^-/-^* than wt compartment of the same chimeric animal (Fig. 4K and L). Altogether, these results show that type I IFN augments the IgG2c^+^ and IgG2b^+^ GC B cell response through direct signaling in B cells and that, in contrast to IFN-γ, this effect goes beyond CSR and enhances the overall magnitude of the GC B cell response.

### Absence of B cell intrinsic type I or type II IFN signaling has limited effects on the core GC B cell transcriptional program but results in reduced T-bet expression and altered isotype composition

Given the direct effect of both type I and type II IFN signaling, we next investigated how the respective IFN family influences the global transcriptional program of B cells within established GC. WT and IFN receptor deficient splenic GC B cells were thus sorted eight days after immunization from the same mixed BM chimeric mice described in Figures 3 and 4. mRNA sequencing analysis was preformed to generate datasets comparing gene expression in WT and *Ifnar*^-/-^ GC B cells or WT and *Ifngr*^-/-^ GC B cells, respectively.

To determine if IFN signaling had altered the core GC B cell program, we first compared these sequencing datasets to the transcriptional changes in GC B cell compared to naïve follicular B cells described in Shi *et al. (44)*. We found that the GC B cell transcriptional signature was largely intact in the absence of either type I or type II IFN signaling (Fig. S2A and B). Key GC B cell transcriptional changes, including mRNA encoding core transcription factors (*Bcl6* and *Bach2*), proteins involved in the somatic hypermutation program (*Aicda* and *Polh*), as well as proteins directing migration and localization of the B cells to the GC (*Sipr2*, *Gpr183*), were unaffected by the loss of B cell intrinsic IFN signaling. This suggests that type I IFN or IFN-γ signaling in B cells is not critical for the GC B cell program. We also compared our IFNAR dataset to the early type I IFN induced transcriptional changes in follicular B cells described by Mostavi *et al. (45)*. Out of 71 genes that showed statistically significant differences in expression with at least a 2-fold change after IFN injection, only six were dysregulated in GC B cells at day eight after immunization with OVA and poly I:C Fig. S2C). This suggests that while a part of the IFN signature may be robustly conserved after IFN signaling has stopped a majority of the IFN depended transcriptional changes are transient.

For both the IFNAR and IFNGR dataset, we filtered genes that displayed at least a 2-fold and statistically significant difference in expression between the WT and IFN receptor deficient cells (Fig. 5A and B). By this approach, both cytokine families were found to influence a relatively limited number of genes, with only 87 and 72 genes being affected by type I IFN and IFN-γ signaling, respectively (Fig. 5C). While the type I and type II IFN gene expression signatures have been difficult to separate when assaying peripheral blood during infections or in autoimmunity *(46, 47)*, only seven genes were found to be dysregulated in both the IFNAR and IFNGR datasets. Of these, upregulation of three genes, (*Ctse, Tbx21, and Igh2c*) were dependent on both IFNAR and IFNGR signaling. The downregulation of these genes in IFN receptor deficient B cells was confirmed by qrt-PCR analysis from the same sorted GC B cell populations (Fig. 5D and E). This suggests that B cell intrinsic type I IFN and IFN-γ signaling predominately affect different aspects of the GC B cell response but that the two pathways synergize to control class switching to IgG2c, likely through the induction of *Tbx21 (21)*. In addition to IgG2c, the differential gene expression identified in the two data sets indicated that both type I IFN and IFN-γ signaling in B cells affect the expression of other Ig isotypes. While both may have a small impact on IgG2b expression, we found type I IFN signaling to be associated with increased *Ighg3* mRNA expression. IFN-γ signaling was instead required for optimal *Igha* expression. Thus, while type I IFN and IFN-γ signaling collaboratively support switching to IgG2c, and possibly IgG2b, they may have unique roles in controlling switching to other isotypes.

**Fig 5.**
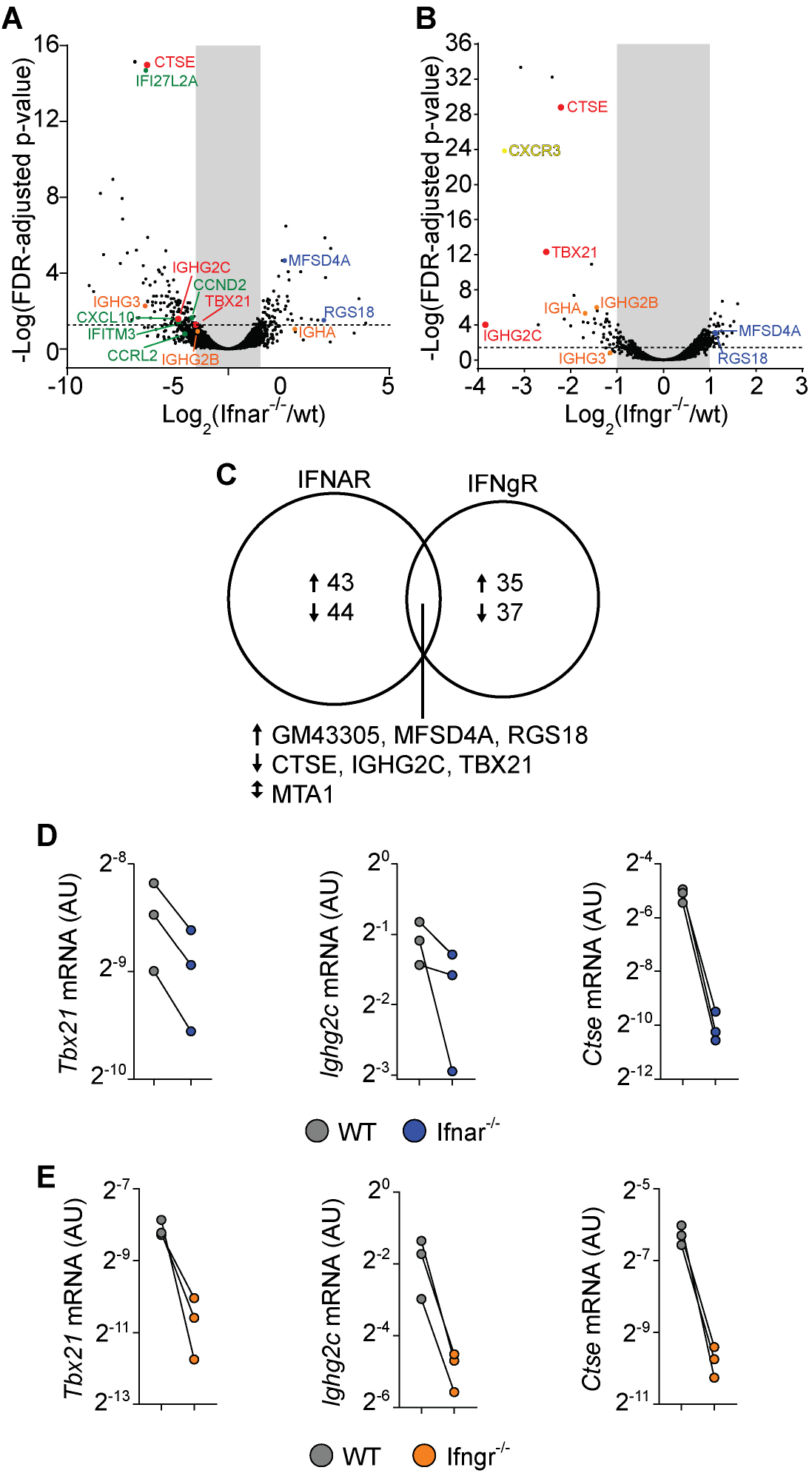
RNA sequencing reveals a limited overlapping effect of B cell intrinsic signaling of Type 1 IFN and IFN-γ on the GC B cell response. Mixed chimeras were generated by reconstituting WT recipients with a 1:1 mix of congenic WT and WT, *Ifnar1^-/-^* or *Ifngr^-/-^* BM cells. 8-10 weeks after reconstitution, chimeras were immunized with OVA + poly I:C, and 8 dpi splenic GC B cells from WT and *Ifnar1^-/-^* **(A)** or *Ifngr1^-/-^* **(B)** were sorted for mRNA sequencing. Volcano plots of changes in gene expression between WT and *Ifnar1^-/-^* **(A)** or *Ifngr1^-/-^* **(B)** GC B cells. **(C)** Venn diagram of genes with 2-fold change and FDR < 0.05 that are regulated by Type 1 IFN or IFN-γ signaling in GC B cells. Analysis of sorted GC B cells by qrt-PCR of genes co-regulated by Type 1 IFNs **(D)** or IFNg **(E)**. Results are from three independent mice.

### IFNAR on non-hematopoietic cells and cognate CD4 T cells is dispensable for Th1 and Tfh cell differentiation after poly I:C adjuvanted immunization

Given that type I IFNs are required for optimal generation of both Tfh and Th1 cells in poly I:C/OVA immunized mice, we next set out to determine the cellular targets for type I IFN signaling in the respective differentiation pathways. To this end we co-transferred CTV-labeled wt and *Ifnar1^-/-^* OT-II cells into BM chimeras, lacking IFNAR in either the hematopoietic, the non-hematopoietic (radio resistant) or both cell compartments. Cell cycle dependent dilution of CTV was examined three days after immunization. The ability of type I IFNs to enhance CD4 T cell proliferation tracked with IFNAR expression by the BM donor cells (i.e. hematopoietic non T cell-intrinsic) (Fig. 6A and B), likely reflecting the ability of type I IFNs to improve the antigen-presenting function and co-stimulatory capacity of APCs *(15, 30, 48, 49)*. No change in CTV dilution was observed when comparing wt and *Ifnar1*^-/-^ OT-II cells or wt and *Ifnar1*^-/-^ irradiated recipient mice, demonstrating that T cell-intrinsic type I IFN signaling or signaling in radio resistant cells has no impact on early T cell proliferation under these conditions (Fig. 6A and B). The ability of type I IFNs to enhance Th1-associated T-bet and IFN-γ (Fig. 6C and D) as well as Tfh cell associated Bcl6 and CXCR5 (Fig. 6E and F) expression was also mostly a result of IFNAR expression on the BM donor cells. However, both T-bet and Bcl6 expression was slightly higher in *Ifnar1*^-/-^ than wt OT-II cells, indicating that type I IFNs to some extent can counteract both Th1 and Tfh cell development through direct effects on T cells (Fig. 6C and E). These effects were however subdominant to the enhanced T-bet and Bcl6 expression driven by T cell extrinsic type I IFN signaling in hematopoietic cells. In conclusion, these results indicate that IFNAR on cognate CD4 T cells or in the non-hematopoietic cell compartment is redundant for both Th1 and Tfh cell fate commitment and hence unlikely to have a major impact on the GC B cell response through these pathways.

**Fig 6.**
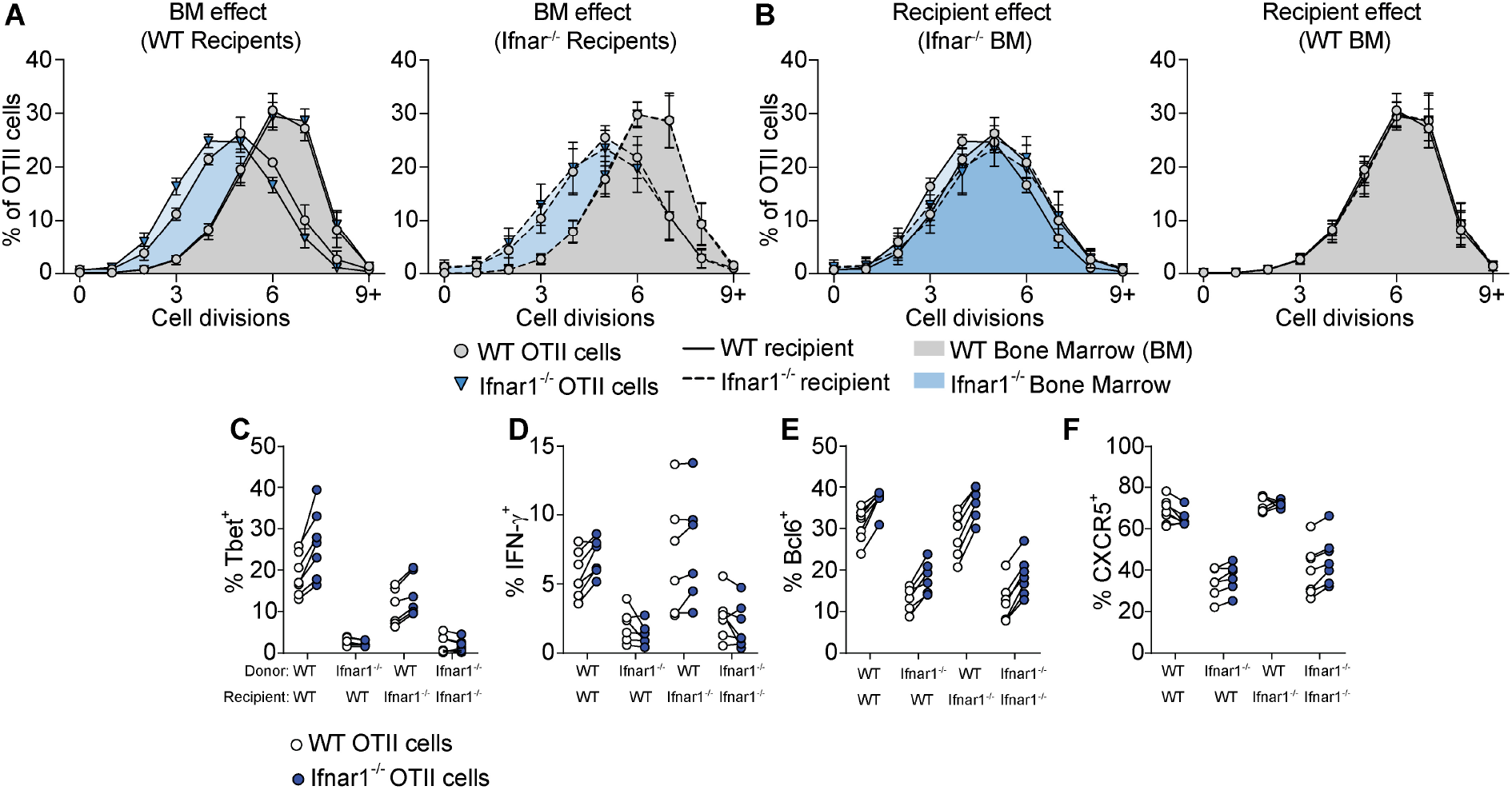
Tfh- and Th1-cell development is supported by IFNAR-signaling in hematopoietic cells distinct from cognate CD4 T cells. Equal numbers (250 000) of CTV-labelled WT and *Ifnar1^-/-^* OT-II cells were co-transferred into BM chimeric recipient mice and spleens were analyzed 3 days after immunization with OVA plus poly I:C. **(A and B)** Percentages (mean ± SD) WT (circle) and *Ifnar1^-/-^* (triangle) OT-II cells in indicated cell cycle number determined by CTV-dilution, broken down by **(A)** recipients or **(B)** donor bone marrow. **(C-F)** Percentage of T-bet^+^ **(C)**, IFN-γ^+^ **(D)**, Bcl6^+^ **(E)**, and CXCR5^+^ **(F)** WT and *Ifnar1^-/-^* OT-II cells, respectively. Results are pooled from two individual experiments consisting of a total of 6-7 mice per group. Each pair of symbols represents one mouse in **(C-F)**.

### Type I IFN signaling in cDCs orchestrates IgG subclass specific GC B cell differentiation through IL-4-secreting Tfh and IFN-γ producing Th1 cells

We have previously demonstrated reduced Tfh cell differentiation in CD11c-cre.*Ifnar1*^fl/fl^ mice with specific deletion of *Ifnar1* in cDCs *(15)*. Here, we wished to determine how conditional deletion of *Ifnar1* in cDCs impacts on the IgG subclass composition within GCs. In particular, while results presented so far reveal how B cell intrinsic type I IFN signaling acts in synergy with the switch factor IFN-γ, produced from cognate CD4 T cells, to enhance IgG2c associated GC B cell responses, it was still unclear why IgG1 ^+^ GC B cells also are strongly reduced in the complete IFNAR knockout (see Fig. 1). Selection of the IgG1 subclass is promoted by IL-4 *(39)* and within secondary lymphoid organs IL-4 secreting T cells are largely confined to the Tfh cell subset *(43, 50)*. To visualize active secretion of IL-4 from cognate CD4 T cells, we intercrossed the OT-II strain with KN2 mice, reporting IL-4 secretion through expression of membrane anchored human CD2 *(51)*. Similar to OT-II cells activated in wild type C57Bl/6 recipients (see Fig. 2D), donor KN2-OT-II cells developed into mutually exclusive Th1 (T-bet^+^) and Tfh (Bcl6^+^) cell subsets eight days after immunization of *Ifnar1^fl/fl^* control mice (hereafter referred to as Cre^-^ control mice) while in *CD11c-Cre.Ifnar1^fl/fl^* mice both subsets were significantly reduced in frequency and number (Fig. 7 A-C). In the Cre^-^ control group, active IL-4 secretion was largely confined to the Bcl6^+^ Tfh cells and although their capacity to produce IL-4 appeared to be at least partially maintained in *CD11c-Cre.Ifnar1^fl/fl^* recipients (Fig. 7D and E), the number of IL-4 secreting Tfh cell was dramatically reduced due to the overall weakened Tfh cell response in these mice (Fig. 7F). Consistent with the impaired generation of T-bet^+^ Th1 cells, IFN-γ producing OT-II cells were also strongly diminished in *CD11c-Cre.Ifnar1^fl/fl^* mice (Fig. 7G). Collectively, these results demonstrate a bifunctional effect of type I IFN signaling in cDCs to promote the appearance of separable IFN-γ producing Th1 and IL-4 producing Tfh cell subsets.

**Fig 7.**
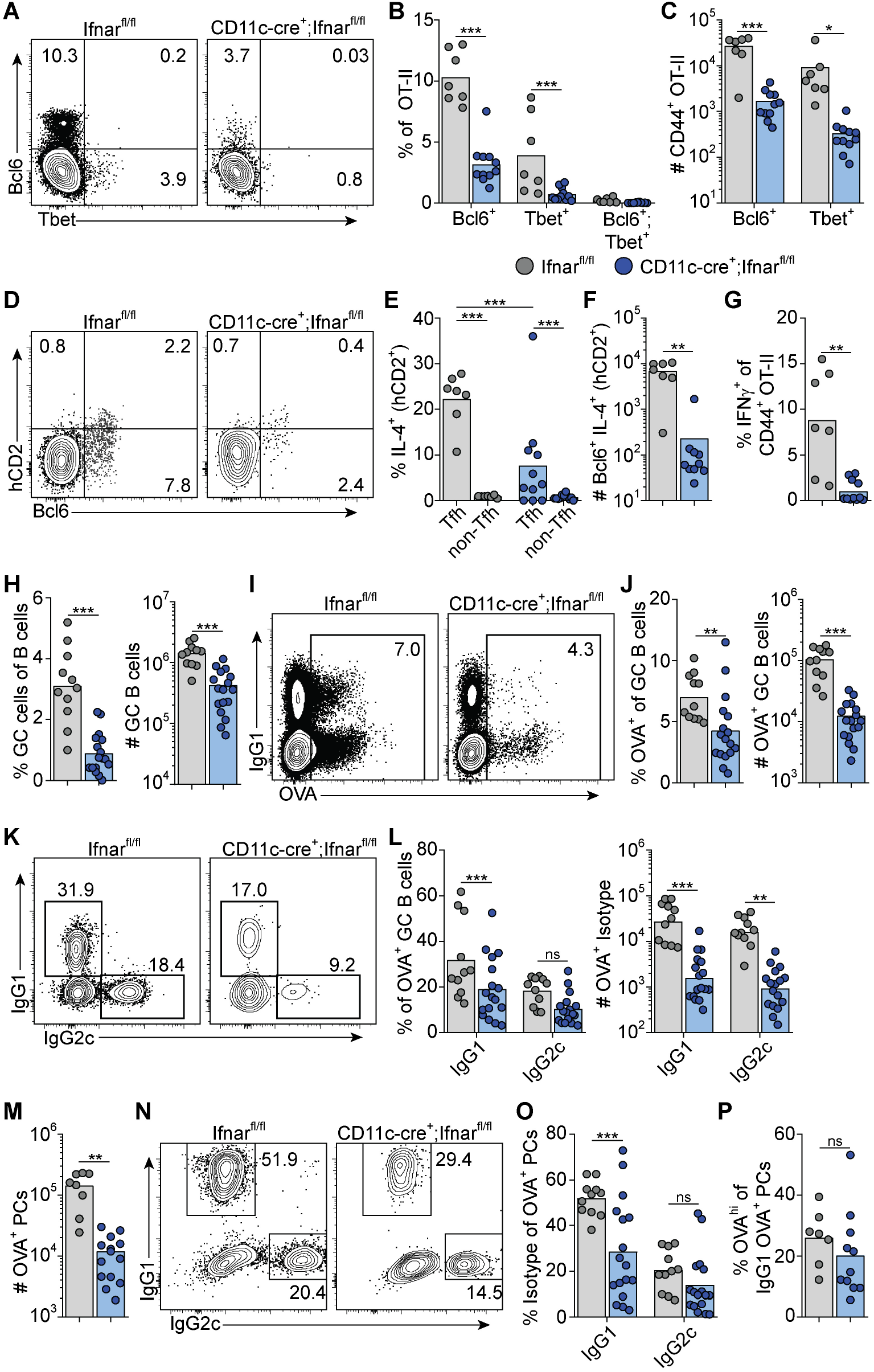
Type 1 IFN signaling in cDC regulates Th1, Tfh and GC B cell development. *Ifnar1^fl/fl^* and *CD11c-cre;Ifnar1^fl/fl^* mice were transferred with 50 000 KN2-OT-II cells and immunized with OVA plus poly I:C and splenocytes were analyzed 8 dpi. **(A)** Flow plots of Bcl6 and T-bet expression in transferred KN2-OT-II cells. Frequency **(B)** and number **(C)** of Bcl6^+^ and T-bet^+^ KN2-OT-II cells. **(D)** Flow plots of hCD2 staining in KN2-OT-II cells. **(E)** Frequency of hCD2^+^ cells in Tfh and non-Tfh KN2-OT-II cells. **(F)** Number of Bcl6^+^ hCD2^+^ KN2-OT-II cells. **(G)** Frequency of IFNg^+^ cells of total KN2-OT-II cells. **(H)** Frequency and number of GC B cells. **(I)** Flow plots of OVA staining of GC B cells. **(J)** Frequency and number of OVA^+^ cells in GC B cells. **(K)** Flow plots of IgG1 and IgG2c expression by OVA^+^ GC B cells. **(L)** Frequency and number of IgG1 and IgG2c cells in OVA^+^ GC B cells. **(M)** Number of OVA^+^ PC. **(N)** Flow plots of IgG1 and IgG2c expression by OVA^+^ PC. **(O)** Frequency of IgG1 and IgG2c cells among OVA^+^ PC. **(P)** Frequency of OVA^hi^ cells within OVA^+^ IgG1^+^ PC. Results are pooled from three independent experiments. Each symbol represents one mouse. *p≤0.05, **p<0.01 and **p<0.001.

Similar to the complete *Ifnar1* knockout, the expansion of GC B cells was strongly reduced after immunization of *CD11c-Cre.Ifnar1^fl/fl^* mice compared with their Cre^-^ controls (Fig. 7H). The requirement for IFNAR-expressing cDC was equally evident when analyzing the number of OVA-binding GC B cells in the two groups of mice, confirming that the reduction in GC B cells was related to the immunization rather than differences in pre-existing GC B cell numbers (Fig. 7I and J). In marked contrast to when B cells specifically lack IFNAR, the reduced GC B cell response in the absence of IFNAR expressing cDCs was not caused by a selective loss of IgG2 expressing GC B cells; IgG1^+^ GC B cells were now at least equally affected, again confirmed by analysis of OVA-binding GC B cells (Fig. 7K and L).

To determine if IFNAR signaling in cDC1 was required for any of the effects that type I IFNs have on Th1, Tfh and GC B cell differentiation, we immunized *XCR1-cre.Ifnar1^fl/fl^* mice *(52)* as above and examined the OT-II cell response (Fig. S3A and B). No difference in OT-II cell expansion or differentiation to Th1 and Tfh cells was observed at day 8 post-immunization. Similarly, no effects on GC B cell expansion or generation of OVA-specific IgG1^+^ or IgG2c^+^ GC B cells were observed (Fig. S3C and D).

Finally, we examined PC generation in immunized *CD11c-Cre.Ifnar1^fl/fl^* mice. Similar to the GC B cell response in these mice we found reduced numbers of OVA-specific PC at day eight post-immunization (Fig. 7M). While the total number of OVA-binding PC thus was more than 10-fold lower in *CD11c-Cre.Ifnar1^fl/fl^* mice as compared with Cre^-^ controls, the reduction was most pronounced for the IgG1 producing subset, as evident from a significant reduction in the percentage of IgG1 ^+^ but not IgG2c^+^ events when comparing the OVA-specific PC that had developed in the two mouse strains (Fig. 7N and O). Of note, the reduced affinity of PCs developing in the complete IFNAR knockout was not apparent in the absence of IFNAR on cDCs, indicating that type I IFN signaling in B cells may underlie this effect (Fig. 7P). Altogether these results show that expansion of IL-4 secreting Tfh cells, through type I IFN signaling in cDCs, represents a third pathway whereby type I IFNs regulate the GC response, providing an explanation to how type I IFN in addition to its pronounced effect on IgG2 also amplifies the IgG1 response.

## Discussion

Type I IFNs possess a wide range of immune stimulatory properties with relevance for antiviral immunity, vaccination, and systemic autoimmune disease. Here, we have dissected how a succinct type I IFN response induced by dsRNA drives GC formation and IgG subclass specification. By selectively blocking receptor signaling in defined target populations or permitting signaling to occur only during defined time windows, we demonstrate that type I IFNs primarily act on B cells and cDCs to rapidly program the B and T cell responses underlying formation of GCs and class switching to a broad IgG subclass distribution. In addition to enhancing generation of Tfh cells and long-lived IgG1 responses, type I IFNs induce T-bet expression and support IgG2c^+^ GC B cell development by mechanisms involving both direct effects on B cells and induction of IFN-γ production by cognate CD4 T cells.

Our results show that type I IFNs act on cDCs within the first 24 hours after immunization to initiate concurrent Tfh and Th1 cell differentiation. Signaling in the cDC1 subset was redundant for both processes. It therefore seems likely that type I IFN signaling in cDC2 plays an important role for both Th cell fates, although we have not addressed this directly through cDC2 specific deletion of IFNAR. The demonstration that type I IFNs enhance both GC B cell responses and Th1 immunity stands in marked contrast to their immune suppressive role in chronic viral infections *(4)*. The LCMV clone 13 strain establishes a persistent infection where long-lasting expression of IFN-β and IFN-a in cDCs leads to exhaustion of the protective Th1 cell response by a mechanism involving up-regulation of PD-L1 and IL-10 expression by the cDCs *(53, 54)*. Likewise, the Th1-promoting property of cDC2 is suppressed by an excessive type I IFN response during severe blood-stage *Plasmodium* infection *(55)*. The stimulatory effects of type I IFNs described in the current study are hence likely related to the short duration of the type I IFN response induced by poly I:C, a notion further supported by the beneficial effects these cytokines have in acute viral infections, including vesicular stomatitis virus (VSV) *(18, 56)*, influenza *(17)*, RSV *(57)* and adenovirus *(12)*. Still, the ability of type I IFNs to concomitantly drive Th1 and Tfh cell development is not necessarily recapitulated in the acute viral infection models. While VSV similar to poly I:C gives rise to a rapid and transient peak of type I IFNs that stimulates generation of Tfh cells, this occurs at the expense of Th1 cell polarization *(16)*. One explanation for this apparent discrepancy could be the mixed pro-inflammatory response induced by viral sensing through multiple innate receptors *(58)*. The adjuvant effect of poly I:C is on the other hand completely lost in IFNAR^-/-^ mice, as shown herein, and injection of purified IFN-β replicates the immune stimulatory effects of poly I:C when co-injected with a protein antigen *(24)*. The induction of T-bet associated GCs and long-lived IgG responses through the isolated effect of type I IFNs should be of considerable interest for the development of adjuvants to enhance antiviral vaccine efficacy. IgG2a/c represents the dominant anti-viral IgG subclass in mice *(59)* and B cell-specific T-bet deletion leads to impaired antiviral IgG2a/c production and viral clearance *(60)*, as well as in an inability to contain chronic viral infection *(61)*. In addition, protective antibody responses against influenza were recently demonstrated to rely on T-bet associated GCs *(23)*. The current study may also contribute towards understanding the immunogenicity of emerging mRNA vaccine approaches *(62)*. Similar to poly I:C, mRNA vaccines induce a short-lived type I IFN response *(32)*, and currently two of twelve SARS-CoV-2 vaccine candidates in phase 3 trials are mRNA-based *(63)* and induce specific IgG serum concentrations higher or equivalent to the levels detected in convalescent sera *(64, 65)*.

While antiviral immunity frequently has been associated with both type I IFN and IgG2a/c antibody production, it has not been clear how type I IFNs promote IgG2a/c dominated GCs. B cell-intrinsic type I IFN signaling was recently shown to be crucial for spontaneous development of IgG2c^+^ GCs in lupus prone *B6.Sle1b* mice *(14)*. GC formation was however not affected by lack of type I IFN signaling when the same lupus-prone strain or wt mice were immunized with NP-conjugated chicken γ-globulin *(14)*, probably reflecting an insufficient type I IFN response to this particular immunization regimen. In the current study, we show that IFN-γ production from cognate CD4 T cells, which requires type I IFN signaling in cDCs, is a critical component of the type I IFN-dependent IgG2c^+^ GC B cell response. This pathway was however not sufficient but acted in concert with direct sensing of type I IFNs by the B cells and both pathways contributed to the induction of T-bet expression in GC B cells. Yet, the effects of type I IFNs and IFN-γ differed. While IFN-γ acted as a non-redundant IgG2c switch factor, only signaling through IFNAR amplified GC B cell expansion with specific effects on IgG2c^+^ and IgG2b^+^ GC B cells. How this occurs and why the effect was confined to GC B cells expressing the IgG2 subclasses remains to be determined. However only early, and not late, IFNAR neutralization resulted in reduced IgG2c^+^ GC B cell numbers. Likewise, only few of the roughly 70 genes previously shown to be induced in B cells following type I IFN treatment *(45)* were affected in IFNAR deficient GC B cells eight days after immunization, further supporting that type I IFNs acted on the B cells early in the response, possibly during initial B cell activation. Indeed, type I IFNs have been shown to confer increased sensitivity to BCR stimulation and to promote B cell expansion by both enhancing proliferation and reducing sensitivity to apoptosis *(14, 66)*.

Whereas targeted deletion of IFNγR in B cells, IFN-γ in cognate CD4 T cells or IFNAR in cDCs resulted in similar reduction in IgG2c^+^ GC B cells, IFNAR deletion in cDCs additionally impaired development of IgG1^+^ GC B cells and resulted in an overall reduced magnitude of the GC response. This coincided with reduced Tfh cell development, with a particularly pronounced effect on IL-4 producing Tfh cells. Tfh cells thus became IL-4 producers also under the strong Th1 polarizing conditions otherwise conferred by poly I:C and downstream type I IFN production. In addition, T-bet was absent from the Tfh cells around the peak of the GC reaction, indicating limited Th1 cell characteristics of the Tfh cell subset. Consistent with this, we demonstrate that both IFN-γ and type I IFN acted on B cells very early after immunization and induced IgG2c CSR prior to evident GC formation. These results are in agreement with previous studies, demonstrating that CSR mostly precedes GC formation *(67–69)*. Nonetheless, similar to viral infection models *(70–72)*, Bcl6 and T-bet were co-expressed by the T cells at this early stage. We could hence not identify divergent Th1 versus Tfh cell commitment at the time when B cells were receiving the IgG2c switch signals. Tfh cells with a history of T-bet expression have been shown to produce IFN-γ within established GCs *(73)*. While it remains possible that equivalent IFN-γ producing T-bet^-^ Tfh cells were present within the GCs studied herein, late IFN-γ neutralization had no detectable effect on the magnitude or IgG subclass composition of the GC response. This indicates that the IgG2c^+^ GC B cells did not rely on continuous IFN-γ signaling within the GCs.

In summary, the current study describes how type I IFNs through at least three separate pathways can enhance and modulate the GC B cell response. Our results provide a detailed roadmap of how this family of cytokines confers long-lived humoral immunity. Exploiting the type I IFN dependent pathways identified herein could provide a means to enhance efficacy of e.g. mRNA vaccine regimens and to prolong the duration of vaccine-induced protection. On the other hand, the relative contribution of these pathways to onset of systemic autoimmune disease warrants further investigations.

## Material & Methods

### Study design

This study sought to determine pleiotropic effects and identify the cellular targets of acute type I IFN signaling on the germinal center B cell response. By selectively blocking receptor signaling in defined target populations, or permitting signaling to occur only during defined time windows, we unravel how GCs with broad IgG subclass distribution develop in mice immunized with OVA plus poly I:C. All experiments were performed according to protocols approved by the Lund/Malmo animal ethical committee (Sweden).

### Mice

C57Bl/6 mice (wild type [wt]) were purchased from Taconic (Ejby, Denmark), *Ifngr1^-/-^* (B6.129S7-*Ifngr1^tm1agt^/J*) and *Ifng^-/-^ (B6.129S7-Ifng^tm1Ts^/J*) mice purchased from The Jackson Laboratory (Bar Harbor, ME, USA). *Il27ra^-/-^* (B6N.129P2-Il27ra^tm1Mak^/J), *Ifnar1^-/-^* (on a C57Bl/6 background), B6.SJL (B6.SJL-Ptprca Pepcb/BoyJ) and OT-II (B6.Cg-Tg(TcraTcrb)425Cbn/J), KN2 (Il4^tm1(CD2)Mmrs^) *(51), CD11c-cre* (B6.Cg-Tg(Itgax-cre)1-1Reiz/J), *XCR1-cre* (Xcr1^Cre-mTFP1^) *(52)* and *Ifnar^fl/fl^* (Ifnar1^tm1Uka^) *(74)* mice were bred and maintained at the Biomedical Center animal facility, Lund University. *Ifnar^fl/fl^* (Ifnar1^tm1Uka^). CD45.1^+^CD45.2^+^OT-II and C57Bl/6 mice were generated by breeding B6.SJL (CD45.1^+^) mice with OT-II or C57Bl/6 (CD45.2^+^) mice, respectively. *Ifng^-/-^* and *Ifnar^-/-^* mice were crossed to OT-II B6xB6.SJL mice to generate *Ifng^-/-^* and *Ifnar^-/-^* OT-II mice. KN2-OT-II mice were generated by crossing KN2 and OTII mice. *CD11c-Cre.Ifnar1^fl/fl^* and *XCR1-cre.Ifnar1^fl/fl^* mice were generated by crossing *Ifnar^fl/fl^* to *CD11c-cre* and *XCR1-cre*, respectively. Mice were included in experiments at 8-12 weeks of age.

### Adoptive transfers, immunizations and mAb treatment

CD4^+^ OT-II cells were isolated from spleen and LNs from OT-II*B6SJL mice with the EasySep mouse CD4^+^ T cell isolation kit (Stemcell Technologies, Vancouver, BC, Canada), according to manufacturer’s protocol. Enriched CD4^+^ cells (>90% purity) were labelled with 5μM CellTrace Violet (Life Technologies, Carlsbad, CA, USA) and 5000 - 5 x 10^5^ cells/recipient were transferred intravenously (i.v.) as indicated. 16-20 hours after transfer, recipients were immunized with 100 μg poly I:C (InvivoGen) together with 300μg OVA or NP-OVA (Biosearch Technologies, Novato, CA, USA), as indicated, by intraperitoneal (i.p.) injection. Type I IFN and IFN-γ signaling was blocked by injection of 1 mg of anti-mIFNAR1 (MAR1-5A3) or anti-mIFN-γ (XMG1.2) respectively, at indicated time-points and control mice were treated with equal amounts of mIgG1 (MOPC-21) or rIgG1 (HRPN), all from BioXcell (West Lebanon, NH, USA).

### Bone marrow chimeras

To generate mixed BM chimeras, BM cells from age matched (8-12 weeks) wt, *Ifnar1^-/-^, Il27r^-/-^* or Ifngr1^-/-^ donor mice were isolated and re-suspended in sterile PBS. A 1:1 mixture of wt and Ifnar1^-/-^,*Il27r^-/-^* or Ifngr1^-/-^ BM cells (2-3×10^6^ total cells) were transferred into lethally irradiated (900 cGy) recipients (CD45.1^+^CD45.2^+^ B6.SJL x C57Bl/6 or CD45.1^+^B6.SJL). Recipient mice were thereafter kept on Ciprofloxacin for 2 weeks. At 8 weeks after transfer, mice were bled to assess reconstitution by flow cytometry. Whole BM chimeric mice were generated by reconstituting irradiated wt (C57Bl/6) and *Ifnar1^-/-^* mice with wt or *Ifnar1^-/-^* BM, otherwise as described above.

### Abs and reagents

Flow-cytometry analyses were performed with Abs conjugated to FITC, PE, PerCP-Cy5.5, allophycocyanin, eFluor 450, Alexa Fluor 700, PE-Cy7, allophycocyanin-Cy7, Brilliant Violet 605, or biotin. The following Abs were used: anti-B220 (RA3-6B2), anti-CD4 (L3T4), anti-IFN-γ (XMG1.2), anti-GL-7 (GL-7), anti-CD38 (90), anti-T-bet (eBio4B10) (eBioscience, San Diego, CA, USA); anti-CXCR5 (2G8), anti-CD62L (MEL-14), anti-CD95 (Jo2), anti-Bcl6 (K112-91) (BD Biosciences, San Jose, CA, USA); anti-IgD (11-26c.2a), anti-CD138 (281-2), anti-CD45.1 (A20), anti-CD45.2 (104), anti-IgM (RMM-1), anti-IgG1 (RMG1-1), anti-IgG2b (RMG2b-1) (BioLegend, San Diego, CA, USA); anti-IgG2c (polyclonal) (Southern Biotech, Birmingham, AL, USA); and donkey anti-rat F(ab’)2 fragment (polyclonal) (Jackson Immunoresearch, West Grove, PA, USA). Streptavidin conjugated to eFluor450 (eBioscience), allophycocyanin (Biolegend), and PE (Southern Biotech) were used as secondary reagents in combination with biotinylated Abs. For detection of NP- or OVA-binding cells, PE-conjugated NP (Biosearch Technologies) or Alexa 647-conjugated OVA (Molecular Probes, Eugene, OR, USA) was used, respectively. Dead cells were excluded using propidium iodide or Live/Dead Fixable Aqua Dead Cell Stain Kit (Molecular Probes).

### Flow cytometry

Single cell suspensions were prepared by mechanical disruption and filtered through 70μm cell strainers. RBCs were lysed with ACK buffer. For IgG analysis, cells were blocked with anti-FcR mAb (2.4G2) in 10% rat serum and thereafter incubated with isotype specific anti-IgG antibodies (see Abs and reagents). Remaining anti-mouse IgG reactivity were subsequently blocked with 10% mouse serum before incubation with fluorophore-conjugated mAbs. CXCR5 was detected as previously described *(15)* and followed by intracellular staining of Bcl6 and T-bet. Intracellular IFN-γ was detected after re-stimulation in complete medium with PMA (50 ng/ml) ionomycin (500 ng/ml; both Sigma-Aldrich, St. Louis, MN, USA), and Brefeldin A (eBioscience) for 3 hours. All intracellular staining was done using the FoxP3 Fixation/Permeabilization kit (eBioscience). Data was acquired on a LSRII or FACS AriaII and analyzed with FlowJo software (BD Bioscience).

### NP-specific ELISA

NP-specific serum antibodies were measured by ELISA. 96-well EIA/RIA plates (Sigma Aldrich) were coated overnight with 0.5ug/ml NP23-BSA in PBS at room temperature and thereafter washed once in wash buffer (PBS+ 0.1% Tween20) and blocked with sample buffer (1% BSA in PBS) for 1 h and thereafter washed twice. Samples were diluted in sample buffer at 1:100 and 1:5000 and subsequently added in duplicates and incubated for 2 hours. Plates were washed four times before biotinylated anti-mouse IgG, anti-mouse IgG2c (polyclonal; both Southern Biotech), or anti-mouse IgG1 (RMG1-1) (BioLegend) was added in sample buffer, incubated for 1 h, and subsequently washed four times. Thereafter, Horseradish peroxidase (HRP) conjugated streptavidin was added and incubated for 45 minutes. After four washes, plates were developed with 3,3’, 5,5’-Tetramethylbenzidine (TMB) and absorbance values were read at 450 nm on a SPECTROstar Nano (BMG Labtech, Ortenberg. Germany). NP-specific standard was prepared by pooling day 14 sera from NP-OVA immunized WT mice and used for all ELISAs to calculate serum titers (determined as 4x background OD value).

### Cell sort and cDNA preparation and quantitative real-time PCR

Total OT-II cells (Live, singlet, CD4^+^ CD45.1^+^ B220^-^) or GC B cells (Live, singlet, B220^+^Fas^+^CD38^-^ CD45.1^+^ or CD45.1^-^) were sorted from immunized recipients directly into RLT buffer supplemented with 1% β-ME (Qiagen, Hilden, Germany). mRNA was extracted with an RNeasy Micro Kit (Qiagen) and either used for sequencing or converted into cDNA using a SuperScript III First-Strand cDNA kit according to manufacturer’s protocol (Thermo Fisher, Waltham, MA, USA). Quantitative RT-PCRs were performed using SYBR GreenER qPCR SuperMix (mo Fisher) with 0.5 μM forward and reverse primers in a final volume of 20 μl. Reactions were run and recorded on an iCycler (Bio-Rad, Hercules, CA, USA).

### Primer sequences

*Actb*: forward; 5’-CCACAGCTGAGAGGGAAATC-3’, reverse; 5’-CTTCTCCAGGGAGGAAGAGG-3’, *Bcl6:* forward; 5’-GTACCTGCAGATGGAGCATGT-3’; reverse; 5’-CTCTTCACGGGGAGGTTTAAGT-3’, *Tbx21*: forward; 5’-CAACAACCCCTTTGCCAAAG-3’; reverse; 5’-TCCCCCAAGCAGTTGACAGT-3’, *Ighg2c*: forward; 5’-CAGACCATCACCTGCAATGT-3’; reverse; 5’-CATGGGGGACACTCTTTGAG-3’, *Ctse*: forward; 5’-ATTCTGGAGGTCTCATAACTTGGAC-3’; reverse; 5’-TGCCAAAGTATTCCATATCCAGGTA-3’.

### mRNA sequencing and analysis

Paired-end RNA sequences were processed with AdapterRemoval (v. 2.1.3) (Lindgreen, 2012), setting the minimum quality to 20, minimum length to 25, collapsing reads when possible and removing Nextera adapters. The first seven base pairs were removed them using seqtk (v. 1.0). For each sample, files with collapsed and singleton reads were aligned to the mouse genome (Ensembl GRCm38.84) using HISAT2 (v. 2.0.1) (Kim et al., 2019). Gene expression profiles were generated using HTSeq (v. 0.11.1) (Anders et al., 2015) with default settings. For each group, genes that had >50 raw counts in >2 samples in either the wildtype- or the knockout condition were included in the analysis. Differential expression analyses were performed in R (R Core Team, 2018) using the edgeR package (McCarthy et al., 2012). Normalization factors were calculated using the trimmed mean of M-values (TMM) method. The edgeR general linear model was used for the differential expression analysis. Differentially expressed genes with a false discovery rate of <5% were considered statistically significant.

### Statistical analysis

Data were analyzed with Prism version 6.0 (GraphPad Software). Analysis of statistical significance was done using one-way ANOVA with Kruskal-Wallis multiple comparison test for tree or more groups, or Mann-Whitney U test for two unpaired groups. Differences were considered significant when p ≤ 0.05 (*p≤0.05, **p<0.01and ***p<0.001).

## Acknowledgement

We thank Dr. Ulrich Kalinke and Dr. Bernard Malissen for providing *Ifnar1^fl/fl^* and *XCR1-cre* mice, respectively. We also thank Drs. Jose Maria Gonzalez-Izarzugaza and Kristine Belling for help with analysis of mRNA sequencing results. This work was supported by the Swedish Cancer foundation (Cancerfonden; 18 0324) and the Lundbeck foundation (R155-2014-4184). KL was supported by a Vetenskapsrådet Young Investigator Award (2014-3595), the Ragnar Söderberg Foundation Fellowship in Medicine, a Lundbeck Foundation Research Fellowship (R215-2015-4100), and the Crafoord Foundation (20200733). KN and SB were supported by the Novo Nordisk Foundation (NNF14CC0001).

## Author contributions

BJL, MWD and AWP conceived of the study and designed experiments. MWD and AWP performed experiments and analyzed results. KN and SB analyzed RNA-seq data. BJL secured funding for the study. SB and KL provided essential resources. BJL supervised the project. MWD, AWP and BJL synthesized results and wrote the manuscript with input from co-authors.

Authors declare no competing interests exist.

## References

1. C. G. Vinuesa, I. Sanz, M. C. Cook, Dysregulation of germinal centres in autoimmune disease, Nature Reviews Immunology 9, 845–857 (2009).

2. S. Crotty, Follicular helper CD4 T cells (TFH), Annu. Rev. Immunol. 29, 621–663 (2011).

3. L. B. Ivashkiv, L. T. Donlin, Regulation of type I interferon responses, Nature Reviews Immunology 14, 36–49 (2014).

4. J. R. Teijaro, Type I interferons in viral control and immune regulation, Curr Opin Virol 16, 31–40 (2016).

5. P. Bastard, L. B. Rosen, Q. Zhang, E. Michailidis, H.-H. Hoffmann, Y. Zhang, K. Dorgham, Q. Philippot, J. Rosain, V. Béziat, J. Manry, E. Shaw, L. Haljasmägi, P. Peterson, L. Lorenzo, L. Bizien, S. Trouillet-Assant, K. Dobbs, A. A. de Jesus, A. Belot, A. Kallaste, E. Catherinot, Y. Tandjaoui-Lambiotte, J. Le Pen, G. Kerner, B. Bigio, Y. Seeleuthner, R. Yang, A. Bolze, A. N. Spaan, O. M. Delmonte, M. S. Abers, A. Aiuti, G. Casari, V. Lampasona, L. Piemonti, F. Ciceri, K. Bilguvar, R. P. Lifton, M. Vasse, D. M. Smadja, M. Migaud, J. Hadjadj, B. Terrier, D. Duffy, L. Quintana-Murci, D. van de Beek, L. Roussel, D. C. Vinh, S. G. Tangye, F. Haerynck, D. Dalmau, J. Martinez-Picado, P. Brodin, M. C. Nussenzweig, S. Boisson-Dupuis, C. Rodríguez-Gallego, G. Vogt, T. H. Mogensen, A. J. Oler, J. Gu, P. D. Burbelo, J. Cohen, A. Biondi, L. R. Bettini, M. D’Angio, P. Bonfanti, P. Rossignol, J. Mayaux, F. Rieux-Laucat, E. S. Husebye, F. Fusco, M. V. Ursini, L. Imberti, A. Sottini, S. Paghera, E. Quiros-Roldan, C. Rossi, R. Castagnoli, D. Montagna, A. Licari, G. L. Marseglia, X. Duval, J. Ghosn, HGID Lab§, NIAID-USUHS Immune Response to COVID Group§, COVID Clinicians†, COVID-STORM Clinicians†, Imagine COVID Group†, French COVID Cohort Study Group†, The Milieu Intérieur Consortium§, CoV-Contact Cohort†, Amsterdam UMC Covid-19 Biobank§, COVID Human Genetic Effort†, J. S. Tsang, R. Goldbach-Mansky, K. Kisand, M. S. Lionakis, A. Puel, S.-Y. Zhang, S. M. Holland, G. Gorochov, E. Jouanguy, C. M. Rice, A. Cobat, L. D. Notarangelo, L. Abel, H. C. Su, J.-L. Casanova, Autoantibodies against type I IFNs in patients with life-threatening COVID-19, Science 129, eabd4585–19 (2020).

6. Q. Zhang, P. Bastard, Z. Liu, J. Le Pen, M. Moncada-Velez, J. Chen, M. Ogishi, I. K. D. Sabli, S. Hodeib, C. Korol, J. Rosain, K. Bilguvar, J. Ye, A. Bolze, B. Bigio, R. Yang, A. A. Arias, Q. Zhou, Y. Zhang, F. Onodi, S. Korniotis, L. Karpf, Q. Philippot, M. Chbihi, L. Bonnet-Madin, K. Dorgham, N. Smith, W. M. Schneider, B. S. Razooky, H.-H. Hoffmann, E. Michailidis, L. Moens, J. E. Han, L. Lorenzo, L. Bizien, P. Meade, A.-L. Neehus, A. C. Ugurbil, A. Corneau, G. Kerner, P. Zhang, F. Rapaport, Y. Seeleuthner, J. Manry, C. Masson, Y. Schmitt, A. Schlüter, T. Le Voyer, T. Khan, J. Li, J. Fellay, L. Roussel, M. Shahrooei, M. F. Alosaimi, D. Mansouri, H. Al-Saud, F. Al-Mulla, F. Almourfi, S. Z. Al-Muhsen, F. Alsohime, S. Al Turki, R. Hasanato, D. van de Beek, A. Biondi, L. R. Bettini, M. D’Angio, P. Bonfanti, L. Imberti, A. Sottini, S. Paghera, E. Quiros-Roldan, C. Rossi, A. J. Oler, M. F. Tompkins, C. Alba, I. Vandernoot, J.-C. Goffard, G. Smits, I. Migeotte, F. Haerynck, P. Soler-Palacin, A. Martin-Nalda, R. Colobran, P.-E. Morange, S. Keles, F. Çölkesen, T. Ozcelik, K. K. Yasar, S. Senoglu, Ş. N. Karabela, C. R. Gallego, G. Novelli, S. Hraiech, Y. Tandjaoui-Lambiotte, X. Duval, C. Laouénan, COVID-STORM Clinicians†, COVID Clinicians†, Imagine COVID Group†, French COVID Cohort Study Group†, CoV-Contact Cohort†, Amsterdam UMC Covid-19, Biobank†, COVID Human Genetic Effort†, NIAID-USUHS, TAGC COVID Immunity Group†, A. L. Snow, C. L. Dalgard, J. Milner, D. C. Vinh, T. H. Mogensen, N. Marr, A. N. Spaan, B. Boisson, S. Boisson-Dupuis, J. Bustamante, A. Puel, M. Ciancanelli, I. Meyts, T. Maniatis, V. Soumelis, A. Amara, M. Nussenzweig, A. Garcia-Sastre, F. Krammer, A. Pujol, D. Duffy, R. Lifton, S.-Y. Zhang, G. Gorochov, V. Béziat, E. Jouanguy, V. Sancho-Shimizu, C. M. Rice, L. Abel, L. D. Notarangelo, A. Cobat, H. C. Su, J.-L. Casanova, Inborn errors of type I IFN immunity in patients with lifethreatening COVID-19, Science, eabd4570–23 (2020).

7. A. N. Theofilopoulos, R. Baccala, B. Beutler, D. H. Kono, Type I interferons (alpha/beta) in immunity and autoimmunity, Annu. Rev. Immunol. 23, 307–336 (2005).

8. E. C. Baechler, F. M. Batliwalla, G. Karypis, P. M. Gaffney, W. A. Ortmann, K. J. Espe, K. B. Shark, W. J. Grande, K. M. Hughes, V. Kapur, P. K. Gregersen, T. W. Behrens, Interferon-inducible gene expression signature in peripheral blood cells of patients with severe lupus, PNAS 100, 2610–2615 (2003).

9. L. Bennett, A. K. Palucka, E. Arce, V. Cantrell, J. Borvak, J. Banchereau, V. Pascual, Interferon and granulopoiesis signatures in systemic lupus erythematosus blood, J. Exp. Med. 197, 711–723 (2003).

10. R. Banchereau, S. Hong, B. Cantarel, N. Baldwin, J. Baisch, M. Edens, A.-M. Cepika, P. Acs, J. Turner, E. Anguiano, P. Vinod, S. Khan, G. Obermoser, D. Blankenship, E. Wakeland, L. Nassi, A. Gotte, M. Punaro, Y.-J. Liu, J. Banchereau, J. Rossello-Urgell, T. Wright, V. Pascual, Personalized Immunomonitoring Uncovers Molecular Networks that Stratify Lupus Patients, Cell 165, 551–565 (2016).

11. A. Le Bon, C. Thompson, E. Kamphuis, V. Durand, C. Rossmann, U. Kalinke, D. F. Tough, Cutting edge: enhancement of antibody responses through direct stimulation of B and T cells by type I IFN, J. Immunol. 176, 2074–2078 (2006).

12. J. Zhu, X. Huang, Y. Yang, Type I IFN signaling on both B and CD4 T cells is required for protective antibody response to adenovirus, J. Immunol. 178, 3505–3510 (2007).

13. A. Das, B. A. Heesters, A. Bialas, J. O’Flynn, I. R. Rifkin, J. Ochando, N. Mittereder, G. Carlesso, R. Herbst, M. C. Carroll, Follicular Dendritic Cell Activation by TLR Ligands Promotes Autoreactive B Cell Responses, Immunity 46, 106–119 (2017).

14. P. P. Domeier, S. B. Chodisetti, S. L. Schell, Y. I. Kawasawa, M. J. Fasnacht, C. Soni, Z. S. M. Rahman, B-Cell-Intrinsic Type 1 Interferon Signaling Is Crucial for Loss of Tolerance and the Development of Autoreactive B Cells, Cell Rep 24, 406–418 (2018).

15. H. Cucak, U. Yrlid, B. Reizis, U. Kalinke, B. Johansson-Lindbom, Type I interferon signaling in dendritic cells stimulates the development of lymph-node-resident T follicular helper cells, Immunity 31, 491–501 (2009).

16. M. De Giovanni, V. Cutillo, A. Giladi, E. Sala, C. G. Maganuco, C. Medaglia, P. Di Lucia, E. Bono, C. Cristofani, E. Consolo, L. Giustini, A. Fiore, S. Eickhoff, W. Kastenmüller, I. Amit, M. Kuka, M. Iannacone, Spatiotemporal regulation of type I interferon expression determines the antiviral polarization of CD4+ T cells, Nat. Immunol. 21, 321–330 (2020).

17. E. S. Coro, W. L. W. Chang, N. Baumgarth, Type I IFN receptor signals directly stimulate local B cells early following influenza virus infection, J. Immunol. 176, 4343–4351 (2006).

18. K. Fink, K. S. Lang, N. Manjarrez-Orduno, T. Junt, B. M. Senn, M. Holdener, S. Akira, R. M. Zinkernagel, H. Hengartner, Early type I interferon-mediated signals on B cells specifically enhance antiviral humoral responses, Eur. J. Immunol. 36, 2094–2105 (2006).

19. D. Markine-Goriaynoff, J.-P. Coutelier, Increased efficacy of the immunoglobulin G2a subclass in antibody-mediated protection against lactate dehydrogenase-elevating virus-induced polioencephalomyelitis revealed with switch mutants, J. Virol. 76, 432–435 (2002).

20. C. Haas, B. Ryffel, M. Le Hir, IFN-gamma is essential for the development of autoimmune glomerulonephritis in MRL/Ipr mice, J. Immunol. 158, 5484–5491 (1997).

21. S. L. Peng, S. J. Szabo, L. H. Glimcher, T-bet regulates IgG class switching and pathogenic autoantibody production, PNAS 99, 5545–5550 (2002).

22. N. S. Wang, L. J. McHeyzer-Williams, S. L. Okitsu, T. P. Burris, S. L. Reiner, M. G. McHeyzer-Williams, Divergent transcriptional programming of class-specific B cell memory by T-bet and RORα, Nat. Immunol. 13, 604–611 (2012).

23. J. L. Johnson, R. L. Rosenthal, J. J. Knox, A. Myles, M. S. Naradikian, J. Madej, M. Kostiv, A. M. Rosenfeld, W. Meng, S. R. Christensen, S. E. Hensley, J. Yewdell, D. H. Canaday, J. Zhu, A. B. McDermott, Y. Dori, M. Itkin, E. J. Wherry, N. Pardi, D. Weissman, A. Naji, E. T. L. Prak, M. R. Betts, M. P. Cancro, The Transcription Factor T-bet Resolves Memory B Cell Subsets with Distinct Tissue Distributions and Antibody Specificities in Mice and Humans, Immunity 52, 842–855.e6 (2020).

24. A. Le Bon, G. Schiavoni, G. D’Agostino, I. Gresser, F. Belardelli, D. F. Tough, Type i interferons potently enhance humoral immunity and can promote isotype switching by stimulating dendritic cells in vivo, Immunity 14, 461–470 (2001).

25. F. D. Finkelman, A. Svetic, I. Gresser, C. Snapper, J. Holmes, P. P. Trotta, I. M. Katona, W. C. Gause, Regulation by interferon alpha of immunoglobulin isotype selection and lymphokine production in mice, J. Exp. Med. 174, 1179–1188 (1991).

26. E. Proietti, L. Bracci, S. Puzelli, T. Di Pucchio, P. Sestili, E. De Vincenzi, M. Venditti, I. Capone, I. Seif, E. De Maeyer, D. Tough, I. Donatelli, F. Belardelli, Type I IFN as a natural adjuvant for a protective immune response: lessons from the influenza vaccine model, J. Immunol. 169, 375–383 (2002).

27. C. L. Swanson, T. J. Wilson, P. Strauch, M. Colonna, R. Pelanda, R. M. Torres, Type I IFN enhances follicular B cell contribution to the T cell-independent antibody response, J. Exp. Med. 207, 1485–1500 (2010).

28. F. D. Finkelman, I. M. Katona, T. R. Mosmann, R. L. Coffman, IFN-gamma regulates the isotypes of Ig secreted during in vivo humoral immune responses, J. Immunol. 140, 1022–1027 (1988).

29. V. E. Schijns, B. L. Haagmans, E. O. Rijke, S. Huang, M. Aguet, M. C. Horzinek, IFN-gamma receptor-deficient mice generate antiviral Th1-characteristic cytokine profiles but altered antibody responses, J. Immunol. 153, 2029–2037 (1994).

30. M. P. Longhi, C. Trumpfheller, J. Idoyaga, M. Caskey, I. Matos, C. Kluger, A. M. Salazar, M. Colonna, R. M. Steinman, Dendritic cells require a systemic type I interferon response to mature and induce CD4+ Th1 immunity with poly IC as adjuvant, J. Exp. Med. 206, 1589–1602 (2009).

31. H. Kumar, H. Kumar, S. Koyama, S. Koyama, K. J. Ishii, K. J. Ishii, T. Kawai, T. Kawai, S. Akira, S. Akira, Cutting Edge: Cooperation of IPS-1-and TRIF-Dependent Pathways in Poly IC-Enhanced Antibody Production and Cytotoxic T Cell Responses, J. Immunol. 180, 683–687 (2008).

32. U. Sahin, P. Oehm, E. Derhovanessian, R. A. Jabulowsky, M. Vormehr, M. Gold, D. Maurus, D. Schwarck-Kokarakis, A. N. Kuhn, T. Omokoko, L. M. Kranz, M. Diken, S. Kreiter, H. Haas, S. Attig, R. Rae, K. Cuk, A. Kemmer-Brück, A. Breitkreuz, C. Tolliver, J. Caspar, J. Quinkhardt, L. Hebich, M. Stein, A. Hohberger, I. Vogler, I. Liebig, S. Renken, J. Sikorski, M. Leierer, V. Müller, H. Mitzel-Rink, M. Miederer, C. Huber, S. Grabbe, J. Utikal, A. Pinter, R. Kaufmann, J. C. Hassel, C. Loquai, Ö. Türeci, An RNA vaccine drives immunity in checkpoint-inhibitor-treated melanoma, Nature, 1–24 (2020).

33. S. W. Jackson, H. M. Jacobs, T. Arkatkar, E. M. Dam, N. E. Scharping, N. S. Kolhatkar, B. Hou, J. H. Buckner, D. J. Rawlings, B cell IFN-γ receptor signaling promotes autoimmune germinal centers via cell-intrinsic induction of BCL-6, J. Exp. Med. 158, jem.20151724 (2016).

34. P. P. Domeier, S. B. Chodisetti, C. Soni, S. L. Schell, M. J. Elias, E. B. Wong, T. K. Cooper, D. Kitamura, Z. S. M. Rahman, IFN-γ receptor and STAT1 signaling in B cells are central to spontaneous germinal center formation and autoimmunity, J. Exp. Med. 160, jem.20151722 (2016).

35. Y. Zhang, L. Tech, L. A. George, A. Acs, R. E. Durrett, H. Hess, L. S. K. Walker, D. M. Tarlinton, A. L. Fletcher, A. E. Hauser, K.-M. Toellner, Plasma cell output from germinal centers is regulated by signals from Tfh and stromal cells, J. Exp. Med. 215, 1227–1243 (2018).

36. P. Marrack, J. Kappler, T. Mitchell, Type I interferons keep activated T cells alive, J. Exp. Med. 189, 521–530 (1999).

37. C. Havenar-Daughton, G. A. Kolumam, K. Murali-Krishna, Cutting Edge: The direct action of type I IFN on CD4 T cells is critical for sustaining clonal expansion in response to a viral but not a bacterial infection, J. Immunol. 176, 3315–3319 (2006).

38. S. J. Szabo, S. T. Kim, G. L. Costa, X. Zhang, C. G. Fathman, L. H. Glimcher, A novel transcription factor, T-bet, directs Th1 lineage commitment, Cell 100, 655–669 (2000).

39. C. M. Snapper, W. E. Paul, Interferon-gamma and B cell stimulatory factor-1 reciprocally regulate Ig isotype production, Science 236, 944–947 (1987).

40. T. Yoshimoto, K. Okada, N. Morishima, S. Kamiya, T. Owaki, M. Asakawa, Y. Iwakura, F. Fukai, J. Mizuguchi, Induction of IgG2a class switching in B cells by IL-27, J. Immunol. 173, 2479–2485 (2004).

41. B. Guo, E. Y. Chang, G. Cheng, The type I IFN induction pathway constrains Th17-mediated autoimmune inflammation in mice, J. Clin. Invest. 118, 1680–1690 (2008).

42. J. Pirhonen, J. Sirén, I. Julkunen, S. Matikainen, IFN-alpha regulates Toll-like receptor-mediated IL-27 gene expression in human macrophages, J. Leukoc. Biol. 82, 1185–1192 (2007).

43. R. L. Reinhardt, H.-E. Liang, R. M. Locksley, Cytokine-secreting follicular T cells shape the antibody repertoire, Nat. Immunol. 10, 385–393 (2009).

44. W. Shi, Y. Liao, S. N. Willis, N. Taubenheim, M. Inouye, D. M. Tarlinton, G. K. Smyth, P. D. Hodgkin, S. L. Nutt, L. M. Corcoran, Transcriptional profiling of mouse B cell terminal differentiation defines a signature for antibody-secreting plasma cells, Nat. Immunol. 16, 663–673 (2015).

45. S. Mostafavi, H. Yoshida, D. Moodley, H. LeBoité, K. Rothamel, T. Raj, C. J. Ye, N. Chevrier, S.-Y. Zhang, T. Feng, M. Lee, J.-L. Casanova, J. D. Clark, M. Hegen, J.-B. Telliez, N. Hacohen, P. L. De Jager, A. Regev, D. Mathis, C. Benoist, Immunological Genome Project Consortium, Parsing the Interferon Transcriptional Network and Its Disease Associations, Cell 164, 564–578 (2016).

46. F. J. Barrat, M. K. Crow, L. B. Ivashkiv, Interferon target-gene expression and epigenomic signatures in health and disease, Nat. Immunol. 20, 1574–1583 (2019).

47. R. Banchereau, A.-M. Cepika, J. Banchereau, V. Pascual, Understanding Human Autoimmunity and Autoinflammation Through Transcriptomics, Annu. Rev. Immunol. 35, 337–370 (2017).

48. M. Montoya, G. Schiavoni, F. Mattei, I. Gresser, F. Belardelli, P. Borrow, D. F. Tough, Type I interferons produced by dendritic cells promote their phenotypic and functional activation, Blood 99, 3263–3271 (2002).

49. J. S. Kurche, C. Haluszczak, J. A. McWilliams, P. J. Sanchez, R. M. Kedl, Type I IFN-dependent T cell activation is mediated by IFN-dependent dendritic cell OX40 ligand expression and is independent of T cell IFNR expression, J. Immunol. 188, 585–593 (2012).

50. I. L. King, M. Mohrs, IL-4-producing CD4+ T cells in reactive lymph nodes during helminth infection are T follicular helper cells, J. Exp. Med. 206, 1001–1007 (2009).

51. K. Mohrs, A. E. Wakil, N. Killeen, R. M. Locksley, M. Mohrs, A two-step process for cytokine production revealed by IL-4 dual-reporter mice, Immunity 23, 419–429 (2005).

52. C. Wohn, V. Le Guen, O. Voluzan, F. Fiore, S. Henri, B. Malissen, Absence of MHC class II on cDC1 dendritic cells triggers fatal autoimmunity to a cross-presented self-antigen, Sci Immunol 5, eaba1896 (2020).

53. J. R. Teijaro, C. Ng, A. M. Lee, B. M. Sullivan, K. C. F. Sheehan, M. Welch, R. D. Schreiber, J. C. de la Torre, M. B. A. Oldstone, Persistent LCMV Infection Is Controlled by Blockade of Type I Interferon Signaling, Science 340, 207–211 (2013).

54. E. B. Wilson, D. H. Yamada, H. Elsaesser, J. Herskovitz, J. Deng, G. Cheng, B. J. Aronow, C. L. Karp, D. G. Brooks, Blockade of Chronic Type I Interferon Signaling to Control Persistent LCMV Infection, Science 340, 202–207 (2013).

55. A. Haque, S. E. Best, M. Montes de Oca, K. R. James, A. Ammerdorffer, C. L. Edwards, F. de Labastida Rivera, F. H. Amante, P. T. Bunn, M. Sheel, I. Sebina, M. Koyama, A. Varelias, P. J. Hertzog, U. Kalinke, S. Y. Gun, L. Renia, C. Ruedl, K. P. A. Macdonald, G. R. Hill, C. R. Engwerda, Type I IFN signaling in CD8-DCs impairs Th1-dependent malaria immunity, J. Clin. Invest. (2014), doi:10.1172/JCI70698.

56. U. Muller, U. Steinhoff, L. F. Reis, S. Hemmi, J. Pavlovic, R. M. Zinkernagel, M. Aguet, Functional role of type I and type II interferons in antiviral defense, Science 264, 1918–1921 (1994).

57. A. Remot, D. Descamps, L. Jouneau, D. Laubreton, C. Dubuquoy, S. Bouet, J. Lecardonnel, E. Rebours, A. Petit-Camurdan, S. Riffault, Flt3 ligand improves the innate response to respiratory syncytial virus and limits lung disease upon RSV reexposure in neonate mice, Eur. J. Immunol. 46, 874–884 (2016).

58. S. Akira, S. Uematsu, O. Takeuchi, Pathogen recognition and innate immunity, Cell 124, 783–801 (2006).

59. J. P. Coutelier, J. T. van der Logt, F. W. Heessen, G. Warnier, J. Van Snick, IgG2a restriction of murine antibodies elicited by viral infections, J. Exp. Med. 165, 64–69 (1987).

60. K. Rubtsova, A. V. Rubtsov, L. F. van Dyk, J. W. Kappler, P. Marrack, T-box transcription factor T-bet, a key player in a unique type of B-cell activation essential for effective viral clearance, Proc. Natl. Acad. Sci. U.S.A. 110, E3216–24 (2013).

61. B. E. Barnett, R. P. Staupe, P. M. Odorizzi, O. Palko, V. T. Tomov, A. E. Mahan, B. Gunn, D. Chen, M. A. Paley, G. Alter, S. L. Reiner, G. M. Lauer, J. R. Teijaro, E. J. Wherry, Cutting Edge: B Cell-Intrinsic T-bet Expression Is Required To Control Chronic Viral Infection, J. Immunol 197, 1017–1022 (2016).

62. N. Pardi, M. J. Hogan, F. W. Porter, D. Weissman, mRNA vaccines — a new era in vaccinology, Nature Publishing Group 17, 261–279 (2018).

63. F. Krammer, SARS-CoV-2 vaccines in development, Nature, 1–16 (2020).

64. L. A. Jackson, E. J. Anderson, N. G. Rouphael, P. C. Roberts, M. Makhene, R. N. Coler, M. P. McCullough, J. D. Chappell, M. R. Denison, L. J. Stevens, A. J. Pruijssers, A. McDermott, B. Flach, N. A. Doria-Rose, K. S. Corbett, K. M. Morabito, S. O’Dell, S. D. Schmidt, P. A. Swanson, M. Padilla, J. R. Mascola, K. M. Neuzil, H. Bennett, W. Sun, E. Peters, M. Makowski, J. Albert, K. Cross, W. Buchanan, R. Pikaart-Tautges, J. E. Ledgerwood, B. S. Graham, J. H. Beigel, mRNA-1273 Study Group, An mRNA Vaccine against SARS-CoV-2 -Preliminary Report, N. Engl. J. Med., NEJMoa2022483 (2020).

65. M. J. Mulligan, K. E. Lyke, N. Kitchin, J. Absalon, A. Gurtman, S. Lockhart, K. Neuzil, V. Raabe, R. Bailey, K. A. Swanson, P. Li, K. Koury, W. Kalina, D. Cooper, C. Fontes-Garfias, P.-Y. Shi, A. X. Z. T. X. reci, K. R. Tompkins, E. E. Walsh, R. Frenck, A. R. Falsey, P. R. Dormitzer, W. C. Gruber, U. X. U. A. X. ahin, K. U. Jansen, Phase 1/2 study of COVID-19 RNA vaccine BNT162b1 in adults, Nature, 1–16 (2020).

66. D. Braun, I. Caramalho, J. Demengeot, IFN-alpha/beta enhances BCR-dependent B cell responses, Int. Immunol. 14, 411–419 (2002).

67. K. M. Toellner, A. Gulbranson-Judge, D. R. Taylor, D. M. Sze, I. C. MacLennan, Immunoglobulin switch transcript production in vivo related to the site and time of antigenspecific B cell activation, J. Exp. Med. 183, 2303–2312 (1996).

68. K. M. Toellner, S. A. Luther, D. M. Sze, R. K. Choy, D. R. Taylor, I. C. MacLennan, H. Acha-Orbea, T helper 1 (Th1) and Th2 characteristics start to develop during T cell priming and are associated with an immediate ability to induce immunoglobulin class switching, J. Exp. Med. 187, 1193–1204 (1998).

69. J. A. Roco, L. Mesin, S. C. Binder, C. Nefzger, P. Gonzalez-Figueroa, P. F. Canete, J. Ellyard, Q. Shen, P. A. Robert, J. Cappello, H. Vohra, Y. Zhang, C. R. Nowosad, A. Schiepers, L. M. Corcoran, K.-M. Toellner, J. M. Polo, M. Meyer-Hermann, G. D. Victora, C. G. Vinuesa, Class-Switch Recombination Occurs Infrequently in Germinal Centers, Immunity 51, 337–350.e7 (2019).

70. L. M. Fahey, E. B. Wilson, H. Elsaesser, C. D. Fistonich, D. B. McGavern, D. G. Brooks, Viral persistence redirects CD4 T cell differentiation toward T follicular helper cells, J. Exp. Med. 208, 987–999 (2011).

71. S. Nakayamada, Y. Kanno, H. Takahashi, D. Jankovic, K. T. Lu, T. A. Johnson, H.-W. Sun, G. Vahedi, O. Hakim, R. Handon, P. L. Schwartzberg, G. L. Hager, J. J. O’Shea, Early Th1 Cell Differentiation Is Marked by a Tfh Cell-like Transition, Immunity 35, 919–931 (2011).

72. J. S. Weinstein, B. J. Laidlaw, Y. Lu, J. K. Wang, V. P. Schulz, N. Li, E. I. Herman, S. M. Kaech, P. G. Gallagher, J. Craft, STAT4 and T-bet control follicular helper T cell development in viral infections, J. Exp. Med. 215, 337–355 (2018).

73. D. Fang, K. Cui, K. Mao, G. Hu, R. Li, M. Zheng, N. Riteau, S. L. Reiner, A. Sher, K. Zhao, J. Zhu, Transient T-bet expression functionally specifies a distinct T follicular helper subset, J. Exp. Med. 215, 2705–2714 (2018).

74. E. Kamphuis, T. Junt, Z. Waibler, R. Förster, U. Kalinke, Type I interferons directly regulate lymphocyte recirculation and cause transient blood lymphopenia, Blood 108, 3253–3261 (2006).

