## Supplemental figures for "Type I interferons promote germinal centers through B cell intrinsic signaling and dendritic cell dependent Th1 and Tfh cell lineages"

### Supplementary Materials:

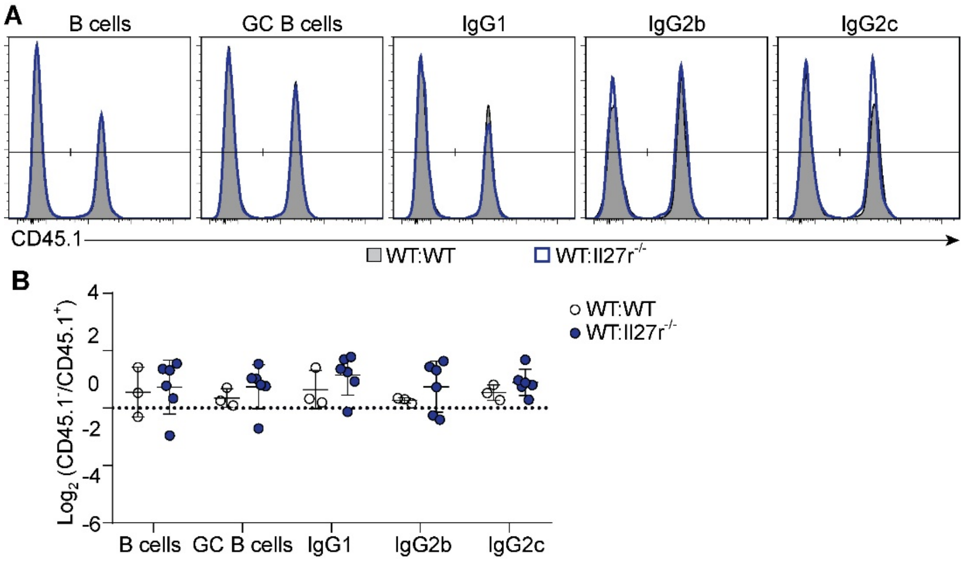


**Fig S1.** IL-27R signaling in B cells is redundant for GC B cell responses. (A-B) Mixed chimeras were generated by reconstituting lethally irradiated WT ($CD45.1^+$, $CD45.2^+$) recipients with a 1:1 mix of WT ($CD45.1^+$, $CD45.2^+$) and WT or *Il27r*$^{-/-}$ ($CD45.1^-$, $CD45.2^+$) BM cells. 8-10 weeks after reconstitution, chimeras were immunized with OVA plus poly I:C, and splenic GC B cell responses were analyzed 8 days later. **(A)** Representative histograms of WT:WT (shaded) and WT:*Il27r*$^{-/-}$ chimeras (blue) showing the distribution of B cells, GC B cells and GC B cells expressing indicated IgG isotypes ($IgG1^+$, $IgG2b^+$ and $IgG2c^+$). **(B)** $Log_2$ normalized ratio of B cells, GC B cells and GC B cells expressing indicated IgG isotype (IgG1, IgG2b and IgG2c) in WT:WT and WT:*Il27r*$^{-/-}$ chimeras. Results are pooled from two (A-B) individual experiments, each symbol represents one mouse.

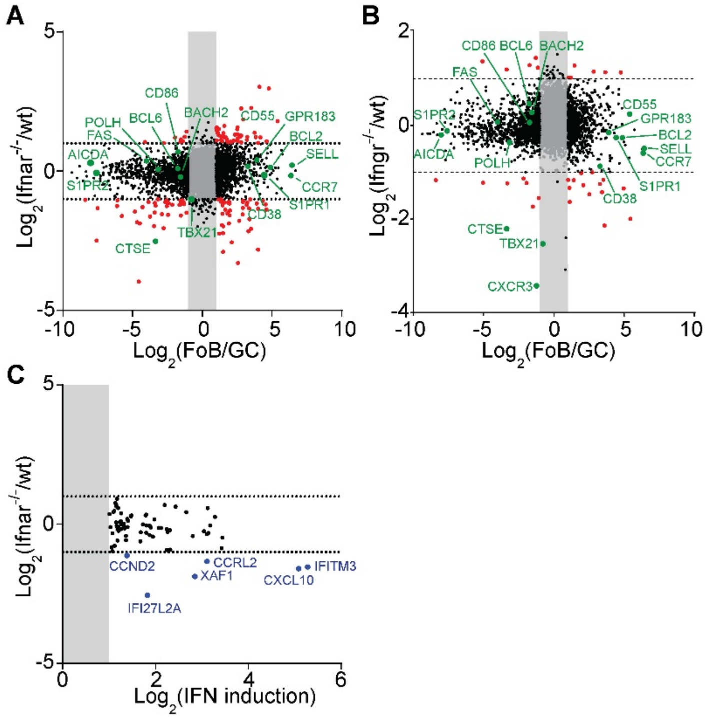


**Fig S2.** The core GC B cell program is largely intact in absence of B cell intrinsic Type 1 IFN and IFN-γ signaling. RNA sequencing data from GC B cells with either Type 1 IFN disruption **(A)** or IFNg **(B)** was compared to sequencing data from Shi *et al*. comparing GC and naïve B cells. **(C)** RNA sequencing data from GC B cells with Type 1 IFN disruption was compared to gene expression changes induced in B cells by injection with Type 1 IFN as described in Mostavi *et al*. Only genes induced by injection of Type 1 IFN with >2-fold change induction and statistical significance were included. Results are from three independent mice.

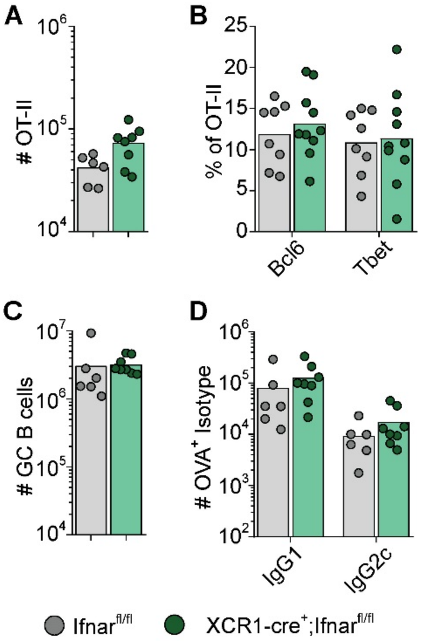


**Fig S3.** Type 1 IFN signaling in cDC1 does not regulate CD4 T cell and GC B cell responses. *Ifnar1fl/fl* and *XCR1-cre;Ifnar1fl/fl* mice were transferred with 50 000 OT-II cells and immunized with OVA plus poly I:C. Lymphocyte responses in the spleen were analyzed 8 days later. **(A)** Number of OT-II cells from *Ifnar1fl/fl* and *XCR1-cre;Ifnar1fl/fl* mice. **(B)** Frequency of Bcl6+ and T-bet+ cells among transferred OT-II. Number of total GC B cell **(C)** and OVA+ GC B cell **(D)** from *Ifnar1fl/fl* and *XCR1-cre;Ifnar1fl/fl* mice. Results are pooled from two independent experiments. Each symbol represents one mouse.
